# Accurate detection of *de novo* and transmitted INDELs within exome-capture data using micro-assembly

**DOI:** 10.1101/001370

**Authors:** Giuseppe Narzisi, Jason A. O’Rawe, Ivan Iossifov, Han Fang, Yoon-ha Lee, Zihua Wang, Yiyang Wu, Gholson J. Lyon, Michael Wigler, Michael C. Schatz

## Abstract

We present a new open-source algorithm, Scalpel, for sensitive and specific discovery of INDELs in exome-capture data. By combining the power of mapping and assembly, Scalpel carefully searches the de Bruijn graph for sequence paths that span each exon. A detailed repeat analysis coupled with a self-tuning *k*-mer strategy allows Scalpel to outperform other state-of-the-art approaches for INDEL discovery. We extensively compared Scalpel with a battery of >10000 simulated and >1000 experimentally validated INDELs against two recent algorithms: GATK HaplotypeCaller and SOAPindel. We report anomalies for these tools to detect INDELs in regions containing near-perfect repeats. We also present a large-scale application of Scalpel for detecting *de novo* and transmitted INDELs in 593 families from the Simons Simplex Collection. Scalpel demonstrates enhanced power to detect long (≥20bp) transmitted events, and strengthens previous reports of enrichment for *de novo* likely gene-disrupting INDELs in autistic children with many new candidate genes.

## Introduction

Enormous advances in next-generation sequencing technologies and computational variation analysis have made it feasible to study human genetics in unprecedented detail. The analysis of Single Nucleotide Variations (SNVs) has become a standard technique and high quality software is available for discovering them [1,2]. However, the same level of performance and reliability is not yet available for detecting *INsertions* and *DELetions* in DNA sequences (INDELs) [3,4]. INDELs are the second most common sources of variation in human genomes and the most common structural variant [5]. Within microsatellites (simple sequence repeats, SSRs, of 1 to 6bp motifs), INDELs alter the length of the repeat motif and have been linked to more than 40 neurological diseases [6]. INDELs also play an important genetic component in autism [7]: *de novo* INDELs that are likely to disrupt the encoded protein are significantly more abundant in affected children than in their unaffected siblings.

Although INDELs play an important role in human genetics, detecting them within next-generation sequencing data remains problematic for several reasons: (1) reads overlapping the INDEL sequence are more difficult to map [8] and may be aligned with multiple mismatches rather than with a gap; (2) irregularity in capture efficiency and non-uniform read distribution increase the number of false positives; (3) increased error rates makes their detection very difficult within microsatellites; and, as shown in this study, (4) localized, near identical repetitive sequences can confound the analysis, creating high rates of false positives. For these reasons, the size of INDELs detectable by available software tools has been relatively small, rarely more than a few dozen base pairs [9].

Two major paradigms are currently used for detecting INDELs. The first and most common approach is to map all of the input reads to the reference genome using one of the available read mappers (e.g., BWA, Bowtie, Novoalign), although the available algorithms are not as effective for mapping across INDELs of more than a few bases. Advanced approaches exploit paired-end information to perform local realignments to detect longer mutations (e.g., GATK UnifiedGenotyper [1, 2] and Dindel [10]), although, in practice, their sensitivity is greatly reduced for longer variants (≥20bp). Split-read methods (e.g., Pindel [11] and Splitread [12]) can theoretically find deletions of any size, but they have limited power to detect insertions due to the short read length of current sequencing technologies (e.g., Illumina). The second paradigm consists of performing *de novo* whole-genome assembly of the reads, and detecting variations between the assembled contigs and the reference genome [13,14]. While having the potential to detect larger mutations, in practice this paradigm is less sensitive: whole-genome sequence assemblers have been designed to reconstruct the high level structure of genomes, while detecting INDELs requires a fine-grained and localized analysis to correctly report homozygous and heterozygous mutations.

Recently, three approaches have been developed that use *de novo* assembly for variation discovery: GATK HaplotypeCaller, SOAPindel [15] and Cortex [16]. The HaplotypeCaller calls SNPs and INDELs simultaneously via local *de-novo* assembly of the haplotypes (http://www.broadinstitute.org/gatk/). Although this is the recommended tool for GATK, the methodological details have not been published and remain unknown. SOAPindel is the variation caller from the Short Oligonucleotide Analysis Package (SOAP). Cortex is a de-novo sequence assembler that uses colored de Bruijn graphs for detecting genetic variants. Cortex was tailored for whole-genome sequencing data and not demonstrated to account for the coverage fluctuations in exome-capture data; thus, it is not used for comparison in this study, although a derivative package, Platypus, is evaluated below. Finally, another recent approach, TIGRA [17], also uses localized assembly, but it has been tailored for breakpoints detection, without reporting the INDEL sequence.

We present a novel DNA sequence micro-assembly pipeline, Scalpel, for detecting INDELs within exome-capture data (Fig. 1). By combining the power of mapping and assembly, Scalpel carefully searches the de Bruijn graph for sequence paths (contigs) that span each exon. The algorithm includes an on-the-fly repeat composition analysis of each exon, coupled with a self-tuning *k*-mer strategy, which allows Scalpel to outperform current state-of-the-art approaches for INDEL discovery specifically in exons containing near-perfect repeat structures.

**Figure 1.**
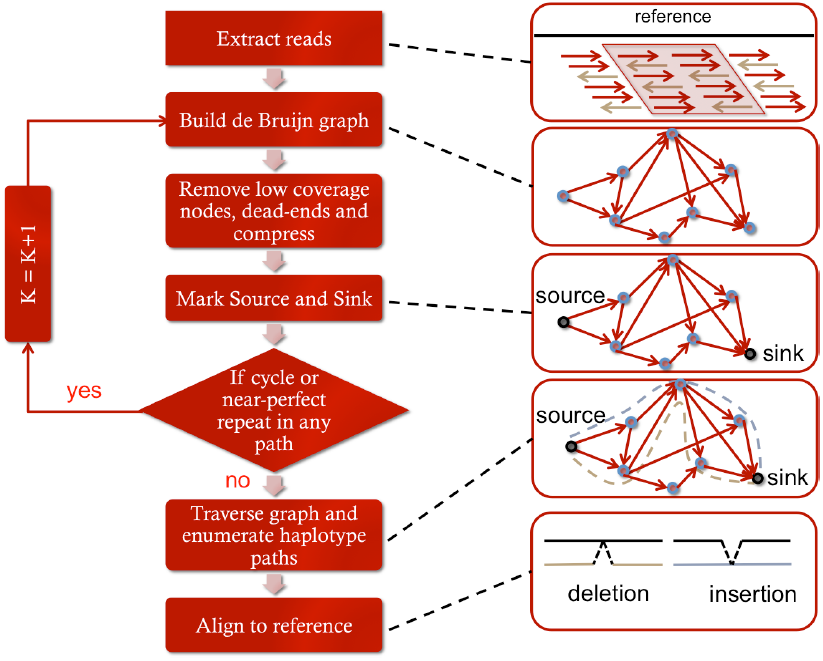
Overview of the Scalpel algorithm workflow. (1) Reads are extracted in the region of interest (e.g., exon) including: (i) well-mapped reads, (ii) soft-clipped reads, and (iii) reads that fail to map, but are anchored by their mate (2) Construct the de Bruijn graph of the input reads. (3) Remove low coverage nodes and dead-ends (sequencing errors), then compress graph. (4) Select source and sink node. (5) Perform on-the-fly analysis of the repeats in each region and increase k-mer if a repeat is found (go to step 2). (6) Source-to-sink graph traversal to find paths that span the region. (7) Align assembled sequences to the reference genome to detect candidate mutations using the standard Smith-Waterman-Gotoh alignment algorithm with affine gap penalties.

## Results

### Experimental validation of variants in one single exome

It is now established that standard mapping methods have reduced power to detect large (≥20bp) INDELs [15, 16] and we confirm this result using simulated reads with 9 algorithms: Scalpel, SOAPindel [15], GATK-HaplotypeCaller, GATK-UnifiedGenotyper, SAMtools [18], FreeBayes [19], Platypus (www.well.ox.ac.uk/platypus), lobSTR [20], and RepeatSeq [21] (**Supplementary Note 1** and **2**). However, the performance of variation discovery tools can change dramatically when applied to real data, so we also performed a large-scale validation experiment involving ∼1000 INDELs from one single exome. The individual has a severe case of Tourette Syndrome and obsessive compulsive disorder (sample ID: K8101-49685), and was sequenced to ≥20x coverage over 80% of the exome target using the Agilent 44MB SureSelect capture protocol and Illumina HiSeq2000 paired-end reads, averaging 90bp in length after trimming. INDELs were called using the three leading pipelines according to their best practices: Scalpel v0.1.1 beta, SOAPindel v2.0.1 and GATK HaplotypeCaller v2.4.3 (Online Methods). Interestingly, there is only ∼37% concordance among calls made by all of the pipelines, and each method reports a variable number of INDELs unique to that pipeline (Fig. 2a). which is in close agreement with the recent analysis [3]. A currently unpublished update to GATK to version 3.0 was released after our initial validation experiments, but we also assessed its accuracy with a second blinded re-sequencing experiment (Fig. 2b and **Supplementary Note 3**).

**Figure 2.**
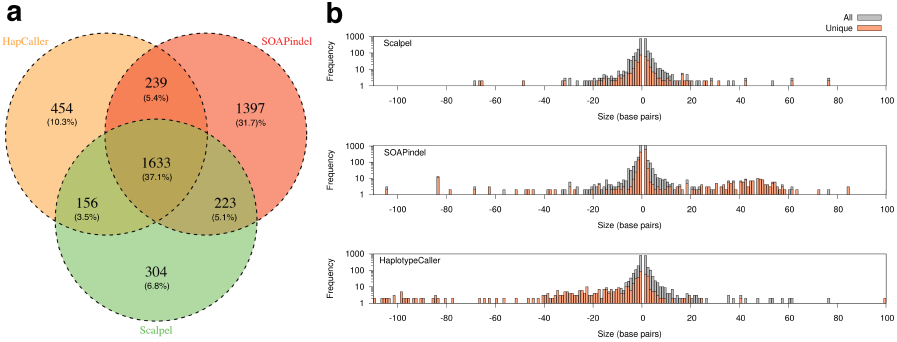
Concordance of INDELs between pipelines. (a) Venn Diagram showing the percentage of INDELs shared between the three pipelines: Scalpel (v0.1.1 beta), SOAPindel (v2.0.1) and GATK HaplotypeCaller (v2.4.3). (b) Size distribution for INDELs called by each pipeline. The whole set of INDELs detected by the pipeline are colored in grey (“All”), while INDELs only called by the pipeline and not by the others are colored in orange (“Unique”).

From the concordance rate alone, it is hard to judge the quality of INDELs unique to each pipeline, as these could either represent superior sensitivity or poor specificity. The size distribution of INDELs called by the HaplotypeCaller (v2.4.3) has a bias towards deletions whereas SOAPindel has a bias towards insertions (Fig. 2b). Scalpel and HaplotypeCaller (v3.0) instead show a well-balanced distribution, in agreement with other studies of human INDEL mutations [9].

We further investigated the performance of the algorithms by a focused re-sequencing of a representative sample of INDELs using the more recent 250bp Illumina MiSeq sequencing protocol (Online Methods). Based on the data depicted in Figure 2a, we selected a total of 1000 INDELs according to the following categories: (1) 200 random INDELs from the intersection of all pipelines; (2) 200 random INDELs only found by HaplotypeCaller (v2.3.4); (3) 200 random INDELs only found by SOAPindel; (4) 200 random INDELs only found by Scalpel; (5) 200 random INDELs of size ≥ 30bp from the union of all three algorithms.

Due to possibly ambiguous representation, INDELs positions are “left-normalized” [3]. However, some ambiguity can still remain, especially within microsatellites, so we computed validation rates using two different approaches. (1) *Position-based:* an INDEL is considered valid if a mutation with the same coordinate exists in the validation data (Fig. 3a). (2) *Exact-match*: an INDEL is considered valid if there is a mutation with the same coordinate and sequence in the validation data (Fig. 3b).

**Figure 3.**
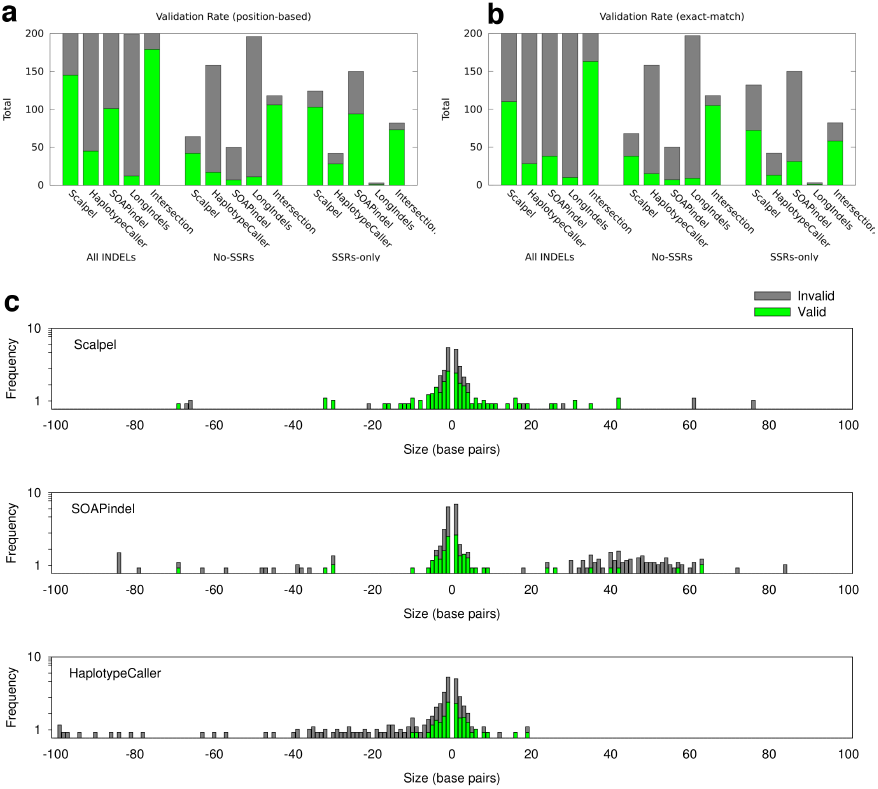
Results of MiSeq validation. (a) Validation rate for different INDEL categories using position-based match. (b) Validation rate for different INDEL categories using exact-match. Results are reported separately for each tool (“Scalpel”, “HaplotypeCaller”, and “SOAPindel”), for all INDELs of size ≥ 30bp from the union of the mutations detected by all three pipelines (“LongIndels”), and for INDELs in the intersection (“Intersection”). Validation results are further organized into three groups: validation for all INDELs (“All INDELs”), validation only for INDELs within microsatellites (“SSRs-only”), and validation for INDELs that are not within microsatellites (“No-SSRs”). (c) Stacked histogram of validation rate by INDEL size for each variant caller. INDELs that passed validation are marked with green color (“Valid”), while INDELs that did not pass validation are marked with grey color (“Invalid”).

As expected, INDELs detected by all pipelines have a high validation rate and their sizes follow a lognormal distribution (**Supplementary Fig. 1**). However, the validation rate varies dramatically for each tool. Respectively, only 22% and 55% of the HaplotypeCaller (2.4.3) and SOAPindel specific INDELs could be validated even when the less strict position-based approach was used, whereas 77% of Scalpel’s specific INDELs are true positive. Even worse is the outcome for the long INDELs: less than 10% passed validation, with SOAPindel and HaplotypeCaller calling the majority of these as erroneous INDELs (Figure 3c and **Supplementary Table 1**). The new version of GATK (v3.0) has largely removed the bias towards deletions (Fig 2b), but find that Scalpel still outperforms HaplotypeCaller (**Supplementary Note 3**). Most significantly, Scalpel shows substantially higher validation rate (76%) for longer INDELs (>5bp) compared to HaplotypeCaller v3.0 (27%).

We further divide the results to separately report the validation rate for INDELs within microsatellites. SOAPindel shows an appreciably higher rate of false-positives within microsatellites (“SSRs-only” in Fig. 3a and 3b). When microsatellites are excluded (“no-SSRs” in Fig. 3a and 3b), the performance of SOAPindel and HaplotypeCaller decline significantly, while Scalpel’s validation rate is only slightly reduced. Figure 3a and Figure 3b also illustrate the relative abundance of INDELs within microsatellites called by each tool, although HaplotypeCaller seems to filter against these. Finally, when switching from position-based to exact-match, INDELs within microsatellites show significant reduction in validation rate. This phenomenon is due to their high instability and higher error rates, and in fact it is not unusual to have more than one candidate mutation at a microsatellite locus.

We further inspected the sequence composition of all false-positive long INDELs. Specifically, we reassembled the 129 SOAPindel invalid long mutations using Scalpel. The majority of these mutations (115) overlap repeat structures where the reference contains a perfect or near-perfect repeat (**Supplementary Fig. 2**). In contrast, of the 62 false-positive long INDELs from HaplotypeCaller, only 16 overlap a repeat. The remaining false positive deletions appear to be due to an aggressive approach used by the algorithm when processing soft-clipped sequences. After manual inspection, the soft-clipped reads in false positive INDELs for HaplotypeCaller are highly variable, and are conjectured to be mapping artifacts of reads from different genomic locations (**Supplementary Fig. 3**). Finally we investigated the relationship between the false-discovery rate (FDR) and characteristic features (e.g., chi-square score and coverage) for 614 INDELs detected by Scalpel and validated by re-sequencing (**Supplementary Note 4**). In addition to highlighting the common trends, this analysis provides recommendation on how to select a chi-square score cutoff to achieve a given FDR.

### Detecting *de novo* and transmitted INDELs in the Simons Simplex Collection

Using Scalpel we detected a total of 3.3 million INDELs in 593 families from the Simons Simplex Collection [22], corresponding to an average of ∼1400 (=3388139/(4*593)) mutations per individual. Accounting for population frequencies, there were 27795 distinct transmitted INDELs across the exomes. Although we detected INDELs only within the exome-capture target regions, we find close agreement to the size distributions reported by Montgomery *et al.* [9] using low coverage whole-genome data from 179 individuals (Figure 4a). Direct comparison to those detected by the GATK-UnifiedGenotyper based mapping pipeline used by Iossifov *et al.* [7] shows that Scalpel has superior power to detect longer insertions (**Supplementary Fig. 4**).

**Figure 4.**
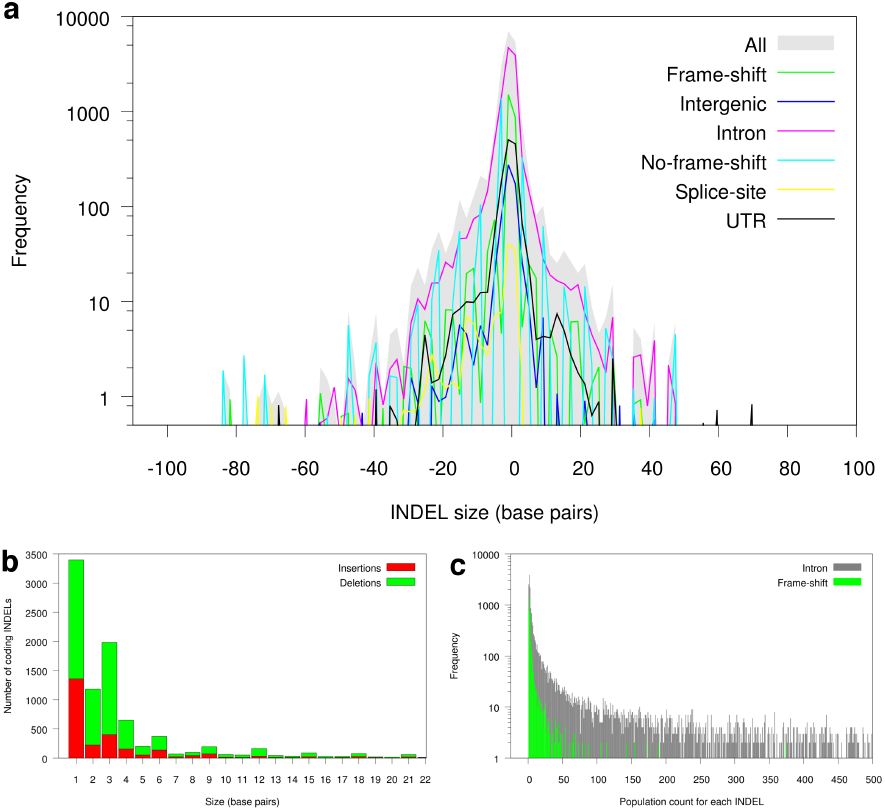
Transmitted mutations in 593 families. (a) Size distribution of insertions and deletions by annotation category. (b) Size distribution of INDELs within coding sequence (CDS). A spike is clearly visible for INDELs with size of multiple of three. (c) Histogram of INDELs frequency by annotation category showing how frame-shifts are typically found at low frequencies in the population.

Despite targeting exons, INDELs are more abundant in introns than other genic locations [5,23] (**Supplementary Fig. 5**). Within the coding sequence (CDS), frame-preserving INDELs are more abundant than frame-shifts (Fig. 4b). In agreement with MacArthur *et al.* [24], we detect a large number of transmitted loss-of-function (LOF) variants in protein-coding genes. Frame-shift mutations are found at lower frequency in the population when located in protein-coding sequences compared to intronic regions (Fig. 4c). Finally, we observe an enrichment of deletions over insertions (**Supplementary Table 2**), with an overall 2:1 ratio across all annotation categories. Similar trends were reported in previous studies [9,23].

To estimate Scalpel’s ability to discover transmitted mutations, we performed targeted re-sequencing of 31 long (≥29bp) transmitted INDELs. Excluding INDELs that failed to sequence (4), 21 passed validation (out of 24), which gives an 87% true positive rate. For three INDELs we could not judge the result of the validation because they were either too long (≥70bp) or they were located in complex regions hard to map.

We had previously reported that *de novo* likely gene disrupting (LGD) mutations, including frame-shift, are significantly more abundant in autistic children than in unaffected siblings by nearly a 2:1 ratio [7]. Other smaller studies came to similar conclusions [25,26,27]. Here we reanalyzed the same data with Scalpel with the goals of confirming such signal, predict additional candidates that could have been missed, and extend the analysis to a larger number of families.

Sanders *et al.* [25] had previously only analyzed SNVs in their study, but our analysis of their 200 families reports the same enrichment for LGD INDELs in autistic children: 11 LGDs in autistic children compared to only 4 in their healthy siblings. In a few cases Scalpel was able to correct the size of the INDEL reported by other algorithms (**Supplementary Note 5** and **6**, **Supplementary Figs. 6-10**). We performed targeted re-sequencing of 102 candidate INDELs; 84 were confirmed as *de novo* mutations, 11 were invalid and 7 failed to sequence, giving an 88% *de novo* positive predictive rate for Scalpel. These reanalysis reveals an important component of deleterious mutations and associated genes that had been previously missed or misreported.

In order to focus the list of candidate genes, we excluded mutations that are common in the population, and used stringent coverage filters (Online Methods) to select a total of 97 high quality *de novo* INDELs (**Supplementary Data 1**). Even after extending the population size from 343 [7] to 593, the same 2:1 enrichment for LGD mutations is confirmed: 35 frame shifts in autistic children vs. 16 in siblings (p-value 0.01097) (**Supplementary Table 3 and 4**). This result also holds for a larger collection of 1303 SSC families (not presented in this study). All together, in agreement with the previously reported results [7], we find a significant overlap between the LGD target genes and the set of 842 FMRP-associated genes [28]. Specifically, 8 out of 35 LGDs in autistic children overlap with the 842 FMRP-associated genes.

Finally we compared the *de novo* variants detected by Scalpel with the ones previously discovered with the GATK-UnifiedGenotyper based mapping pipeline used by Iossifov *et al.* [7] for a larger collection of 716 families sequenced at CSHL (not reported in this study). 21 mutations, out of 175 reported by GATK, were not found by Scalpel. Manual inspection of these loci, indicated that these mutations were below the coverage thresholds used by Scalpel and/or in regions hard to assemble due to the presence of a complex repeat

## Discussion

Assembly is the missing link towards high accuracy and increased power for INDEL mutation discovery for two reasons: (1) it allows the algorithm to break free from the expectations of the reference and (2) extends the power of the method to detect longer mutations. These features are crucial for the analysis of inherited and somatic mutations. Although these ideas have been explored recently in the literature, currently available tools for variant detection suffer from a high error rate. Scalpel is a powerful new method for detecting INDELs in next-generation sequencing data that combines the power of assembly and mapping into a unified framework. While featuring enhanced power to detect longer mutations, Scalpel does not lose specificity. Although we showed results only for exome-capture data, the Scalpel algorithm is agnostic to the sequencing protocol and can be used for whole-genome data as well. We envision that Scalpel will play an important role in the near future for the analysis of inherited and somatic mutation in human studies.

## Acknowledgments

The project was supported in part by National Institutes of Health award (R01-HG006677) and National Science Foundation award (DBI-1350041) to M.C.S., the CSHL Cancer Center Support Grant (5P30CA045508), the Stanley Institute for Cognitive Genomics, and the Simons Foundation (SF51 and SF235988) to M.W. The DNA samples used in this work are included within SSC release 13. Approved researchers can obtain the SSC population dataset described in this study by applying at https://base.sfari.org. We thank S. Eskipehlivan for the technical assistance with the MiSeq validation experiments. We thank M. Bekritsky, S. Neuburgerand, M. Ronemus, D. Levy, B. Yamron, and B. Mishra for helpful discussions and comments on the paper. We thank R. Aboukhalil for testing the software.

## Authors contributions

G.N. developed the software and conducted the computational experiments. G.N. and M.C.S. designed and analyzed the experiments. Y.W. assisted in designing the primers and performed the MiSeq validation experiments. J.A.O. designed the primers and analyzed the MiSeq data. H.F. and J.A.O. assisted with the computational experiments for the comparative analysis between different variant detection pipelines. G.J.L. planned and supervised the experimental design for INDEL validation. Z.W. designed the primers and performed experiments for the validation of *de novo* and transmitted INDELs in the SSC. I.I., Y.L., and M.W. assisted with the analysis of the SSC. G.N. and M.C.S. wrote the manuscript with input from all authors. All of the authors have read and approved the final manuscript.

## Online Methods

### The Scalpel pipeline

Scalpel is designed to perform localized micro-assembly of specific regions of interest in a genome with the goal of detecting insertions and deletions with high accuracy. It is based on the de Bruijn graph assembly paradigm where the reads are decomposed into overlapping *k*-mers, and directed edges are added between *k*-mers that are consecutive within any read [29]. Figure 1 shows the high level structure of the pipeline. (1) The pipeline begins with a fast alignment of the reads to the reference genome using BWA [30,31]. Importantly, these alignments are not directly used to call variations, but only to localize the analysis by identifying all the reads that have similarity to a given locus. Reads are then extracted in the region of interest (e.g., exon) including: (i) *well-mapped* reads, (ii) *soft-clipped* reads, and (iii) reads that *fail to map*, but are anchored by their mate. The latter two classes correspond to locations where the mapper encountered trouble aligning the reads, especially because of the large INDELs present, so it’s necessary to include them in the assembly. (2) Once localized, the algorithm computes an on-the-fly assembly of the reads in the current region using the de Bruijn graph paradigm, specifically, reads are decomposed into overlapping *k*-mers (starting with a default *k*=25) and the associated graph is constructed. (3) Using the reference sequence, one source node and one sink node are selected according to the procedure described later in the “Graph traversal” section. (4) An on-the-fly analysis of the repeats in each region is used to automatically select the *k*-mer size to be used for the assembly, described in section “Repeat analysis”. (5) The graph is then exhaustively examined to find end-to-end paths that span the region. (6) After the sequences are assembled, they are aligned to the reference to detect candidate mutations using a sensitive gapped sequence aligner based on the Smith Waterman algorithm [32] targeted at the reference window. Finally, the above assembly process is applied using a sliding window approach over each target region. By default a window size of 400bp is used with a sliding factor of 100bp. The sliding window strategy is fundamental to handle the highly non-uniform read distribution across the target (**Supplementary Fig. 11**). A window size of 400bp is large enough to assemble the majority of the exons into a single contig: ∼95% of the human exon-targets are shorter than 400bp (**Supplementary Fig. 12**), however each assembly task is small enough for using in-depth techniques to optimize the assembly.

### Graph construction

Two critical components of the Scalpel algorithm are (i) construction of the de Bruijn graph and (ii) detection of sequence paths spanning the targeted region. Reads aligning to the region are extracted and decomposed into overlapping *k*-mers. In order to model the double stranded nature of the DNA, a bidirected de Bruijn graph is constructed [33,34]. The graph is then compressed by merging all non-branching chains of *k*-mers into a single node. Tips and low coverage nodes are removed according to input threshold parameters to remove obvious sequencing errors. Note that, differently from a traditional de Bruijn graph assembler, Scalpel does not use any threading strategy to resolve collapsed repeats. In principle, threading would allows resolution of repeats whose lengths are between *k* and the read length. However we observed in both real and simulated data that, due to the localized graph construction, if a bubble were not covered end-to-end by the reads, threading would either disconnect the graph or introduce errors. Repeats are instead handled differently, as explained in the next section,

### Repeat Analysis

Due to the highly non-uniform read depth distribution across the targeted region and the presence of near-perfect repeats that can mislead the assembly (**Supplementary Note 6** and **Supplementary Fig 13**), Scalpel implements a detailed repeat composition analysis coupled with a self-tuning *k*-mer strategy. Specifically, when assembling a window, Scalpel inspects both the base pair composition of the corresponding reference sequence as well as the resulting de Bruijn graph for the presence of cycles in the graph or near-perfect repeats in the assembled sequences. If a repeat structure is detected, the graph is discarded and a larger *k*-mer is selected. This process continues until a maximum *k*-mer length is reached, which is a function of the read length. If no *k*-mer value can be chosen to avoid the presence of repeats, the region is skipped and the next available region from the sliding window scheme is analyzed. This conservative strategy reduces the number of false-positive calls in highly repetitive regions, and, skips less than 2% of possible windows in the human exome. Note also that, once *k* is selected by the self-tuning k-mer strategy, the graph is “repeat free”, and there is no need to use threading to resolve small repeats.

The proposed self-tuning *k*-mer strategy is similar to the dynamic approach used by SOAPindel and TIGRA to reconnect a broken path in low coverage regions. However, SOAPindel searches for unused reads with gradually shorter *k*-mers until a path is formed or the lower bound on *k*-mer length has been reached; In TIGRA the user can specify the list of *k*-mers to use (by default only two: 15 and 25). Scalpel instead starts from a small *k*-mer value (input parameter) first and then gradually increases it, such that the smallest possible *k*-mer value that generates a “repeat free” graph is used for each region. This strategy has the advantage of better handling of repetitive sequences, highly polymorphic regions, and sequencing errors: source and sink have higher chance to be selected (see section “Graph traversal”) and a smaller *k*-mer reduces the chance of fragmented assembly in low coverage regions.

### Graph traversal

Once a valid de Bruijn graph is constructed, Scalpel examines the graph to find end-to-end sequence paths that span the target window. Because the coverage from exome capture data is highly variable, a special selection algorithm is used to find the edges of each window where coverage is present. First, two nodes in the graph are labeled as *source* and *sink* according to the following procedure: the reference sequence of the target region is scanned left-to-right to detect the first sequence of *k* bases that exactly matches one of the *k*-mers from the nodes in the graph, this node will be marked as the source. In a similar fashion the sink node is detected scanning the reference sequence right-to-left. Since every region is first inspected for repeats, source and sink can be safely selected at this stage. The automated strategy used by Scalpel to select the boundaries of the reference sequence improves over TIGRA’s approach, where the reference region is selected based only on input parameters. After the source and sink nodes are identified, all possible source-to-sink paths are enumerated up to a max number (default 100,000) using a depth-first search (DFS) traversal of the graph, similarly to Sutta assembly algorithm [35]. Note that since the regions to assemble are very small, time and space computational complexities associated with large-scale whole-genome assembly are not relevant and an exact brute-force strategy can be efficiently applied.

If there are no repeat structures in the graph, all the candidate paths are enumerated and aligned to the portion of the reference sequence delimited by source and sink *k*-mers using the standard Smith-Waterman-Gotoh alignment algorithm with affine gap penalties. The list of candidate mutations is then generated. Under typical conditions, the assembler reports a single path for homozygous mutations and two paths for heterozygous mutations. For example, if the sample has an insertion in only one of the two haplotypes, the assembler would discover the INDEL and also the unmodified reference sequence. Note that a traditional sequence assembler would have selected only one of these two paths (usually with higher coverage) and discarded the other one. Scalpel instead examines both paths to distinguish, for example, between homozygous and heterozygous mutations. However, in practice, various factors in real data complicate the detection process and, sometimes, multiple paths are reported in the case of more exotic variations. For example, the Illumina sequencing platform is particularly error prone around microsatellites (e.g., homopolymer runs) and, as a consequence, multiple candidate alleles are elucidated by the data at these loci. Highly polymorphic regions are also prone to generate multiple paths and could be computationally demanding: if the distance between multiple nearby mutations is larger than the (automatically) selected *k*-mer value, each of the associated bubbles in the graph will give rise to two different paths. Finally it is important to note that SNVs are also computed by Scalpel but they are not reported in output. SNVs are used internally for important downstream analysis of the variants called in order, for example, to compute coverage around the indels and to correctly characterize the zygosity of the mutations.

### Exome capture data

Exome capture for the sample K8101-49685 was carried out using the Agilent 44MB SureSelect protocol and then sequenced on Illumina HiSeq2000 with average read length of 100bp. More than 80% of the target region was covered with depth of 20 reads or more. All of the HiSeq data have been submitted to the Sequence Read Archive (http://www.ncbi.nlm.nih.gov/sra) under project accession number SRX265476.

### MiSeq validation

A total of 1,400 INDELs were selected for MiSeq validation over the course of this study. MiSeq validation was initially performed on 1,000 INDELs prior to the release of GATK v3.0 (see the Results section for detailed selection criteria). After the release of GATK v3.0, we selected an additional 400 INDELs for MiSeq validation (see **Supplementary Note 3** for detailed selection criteria). Out of these 400 INDELs, 215 were covered with more than 1,000 reads in the initial MiSeq dataset or in another MiSeq dataset reported by O’Rawe *et al.* [3]. Thus, the second MiSeq validation experiment was performed on the remaining 185 INDELs. For both of the MiSeq validation experiments performed during the course of this study, PCR primers were designed using Primer 3 (http://primer3.sourceforge.net) to produce amplicons ranging in size from 200 to 350 bp, with INDELs of interest located approximately in the center of each amplicon. Primers were obtained from Sigma-Aldrich^®^ in 96-well mixed plate format, 10 μmol/L dilution in Tris per Oligo. Upon arrival, all primers were tested for PCR efficiency using a HAPMAP DNA sample (Catalog ID NA12864, Coriell Institute for Medical Research, Camden, NJ, USA) and LongAmp® Taq DNA Polymerase (New England Biolabs, Beverly, MA, USA). PCR products were visually inspected for amplification efficiency using agarose gel electrophoresis. For the validation experiment, this same PCR was performed using sample K8101-49685 genomic DNA as template. PCR product was verified on E-Gel® 96 gels (Invitrogen Corp., Carlsbad, CA, USA) and subsequently pooled for ExoSAP-IT® (Affymetrix Inc., Santa Clara, CA, USA) cleanup. The cleanup product was further purified using QIAquick PCR Purification Kit (QIAGEN Inc., Valencia, CA, USA) and quantified by Qubit® dsDNA BR Assay Kit (Invitrogen Corp.). Library construction for the MiSeq Personal Sequencer platform (Illumina Inc.) was performed based on Illumina® TruSeq® DNA Sample Prep LS protocol (for the initial 1,000 INDELs) and TruSeq® Nano DNA Sample Preparation Guide (for the additional 185 INDELs), omitting the DNA fragmentation step. Finally, before being loaded onto the MiSeq machine, the quality and quantity of the sample was again verified using the Agilent DNA 1000 Kit on the Agilent Bioanalyzer and with quantitative PCR (Kapa Biosystems Inc., Woburn, MA, USA). This protocol generated high quality 250 bp reads (paired end). The reads were aligned with BWA-MEM (v0.7.5a) to the reference human genome hg19. The alignment was sorted with SAMtools (v0.1.18) and PCR duplicates were marked with the Picard tool set (v1.91). INDELs were realigned with the GATK (version v2.6-4) using the IndelRealigner and base quality scores were recalibrated. Variants were then called with GATK UnifiedGenotyper. All of the MiSeq data have been submitted to the Sequence Read Archive (http://www.ncbi.nlm.nih.gov/sra) under project accession number SRX386284.

### Alignment

Sequencing reads from K8101-49685 exome-capture data were aligned using BWA (v0.6.2-r126) with default parameters to the human reference hg19. Alignments were converted from SAM format to sorted, indexed BAM files with SAMtools (v0.1.18). The Picard tool set (v1.91) was used to remove duplicate reads. These BAM files were used as input for all the INDEL callers used in this study. Reads coming from the re-sequencing experiments were also aligned using BWA. However, if the INDEL approaches half the size of the read length, even after target re-sequencing, mapping the reads containing the INDEL is problematic. The problem is emphasized if the INDEL is located towards the ends of the read (instead of in the middle). To avoid this problem we aligned sequencing reads containing long INDELs (≥30 bp) using Bowtie2 [36] instead of BWA. Bowtie2 offers an end-to-end alignment mode that searches for alignments involving all of the read characters, also called an “untrimmed” or “unclipped” alignment. Specifically, we used the following parameter settings: “--end-to-end --very-sensitive--score-min L,-0.6,-0.6 –rdg 8,1 –rfg 8,1 --mp 20,20”.

### Variant Calling

INDELs for K8101-49685 were called using Scalpel, GATK HaplotypeCaller and SOAPindel as follows:

**Scalpel**. Scalpel (v0.1.1 beta) was run on the indexed BAM using the following parameter setting: “--single --lowcov 1 --mincov 3 –outratio 0.1 --intarget”. INDELs showing high coverage unbalance were then removed (chi-square *k*-mer score > 20).

**GATK**. GATK software tools (v2.4-3 and v3.0) were used for improvement of alignments and genotype calling and refining with recommended parameters. BAM files were re-aligned with the GATK IndelRealigner, and base quality scores were re-calibrated by the GATK base quality recalibration tool. Genotypes were called by the GATK UnifiedGenotyper and HaplotypeCaller. According to the GATK recommendations, the Variant Quality Score Recalibration (VQSR) was not used for the K8101-49685 single exome experiment. Also hard filtering criteria (such us “QD < 2.0, ReadPosRankSum < -20.0 FS > 200.0”) were not used for v2.4.3 as they were aggressively removing long indels from the HaplotypeCaller calls (Supplementary Note 3), but were used instead for the more recent HaplotypeCaller from GATK v3.0. It is important to report that the bias towards deletions for HaplotypeCaller has been extensively reduced with the release of the unpublished GATK 2.8 and 3.0 in December 2013 and March 2014 after our initial MiSeq re-sequencing experiments were completed (Supplementary Note 3). However, many research groups have already employed the older version of HaplotypeCaller for genetic studies and they are still extensively using it. We expect GATK, along with our own and other algorithms, to improve over time as new insights are made into the mutation mechanisms and error profiles.

**SOAPindel.** SOAPindel (v2.0.1) was run on the indexed BAM file using default parameters. According to SOAPindel documentation, putative INDELs are initially assumed to be located near the unmapped reads whose mates mapped to the reference genome. SOAPindel then executes a local assembly (*k*-mer=25 by default) on the clusters of unmapped reads. The assembly results were aligned to reference in order to find the potential INDELs. To distinguish true and false positive INDELs, SOAPindel generates Phred quality scores, which take into consideration the depth of coverage, INDEL size, number of neighboring variants, distance to the edge of contig, and position of the second different base pair. Only those INDELs filtered by internal threshold are retained in the final INDEL call set.

Finally, for all pipelines we selected only INDELs located within the regions targeted by the exome capture protocol.

### The Simons Simplex Collection

The Simons Simplex Collection (SSC) is a permanent repository of genetic samples from 2,700 families operated by SFARI (http://sfari.org) in collaboration with 12 university-affiliated research clinics. Each simplex family has one child affected with autism spectrum disorder, and unaffected siblings. Each genetic sample also has an associated collection of phenotype measurements and assays. The results presented in this work are based on a subset of the SSC composed of 593 families (2372 individuals). Specifically this subset of the SSC collection corresponds to families that have been examined in three recent studies: 343 families from Iossifov *et al.* [7] (CSHL), 200 families from Sanders *et al.* [25] (Yale), and 50 families from O’Roak *et al.* [26] (University of Washington). We selected only family units of four individuals (father, mother, proband, one unaffected sibling), referred to as “quads,” for all analyses in this study.

### Analysis of *de novo* INDELs related to Autism

After eliminating all candidate positions that are common in the population, and thus unlikely to be related to the disorder, we re-assembled each region associated with the candidates INDELs across the family members using a more sensitive parameter setting for Scalpel. Specifically we reduce the starting *k*-mer value to 10 and turned off the removal of low coverage nodes. This step was important to adjust for possible allele imbalance favoring the reference allele over the mutation in the parents, but was impractical to do initially for the whole collection: lowering the *k*-mer and keeping all the nodes in the graph significantly increase the computation complexity of the algorithm. Then we selected de novo INDELs with chi-square *k*-mer score ≤ 10.84.The chi-square *k*-mer score is computed using the standard formula for the chi-square test statistics (χ^2^) but applied to the *k*-mer coverage of the reference and alternative alleles for the mutation according to the following formula:

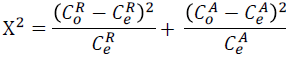

where 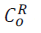 and 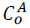 are the observed *k*-mer coverage for the reference and alternative alleles, and 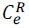 and 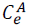 are the expected coverage such that 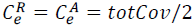. Finally we enforced parents to have at least a *k*-mer coverage of 15 over the assembled region.

### System requirements and software availability

Scalpel is written in C++ and Perl. The source code is freely available as an open-source software project on the SourceForge website at http://scalpel.sourceforge.net. It usually takes 2-3 hours to process one exome-capture data set (80% of target at ≥ 20X) using 10 cores and requiring a minimum of 3GB of RAM.

