## Supplementary Information for "Accurate detection of *de novo* and transmitted INDELs within exome-capture data using micro-assembly" (from another version of this preprint: version 2, 2014-04-15)

#### Supplementary Note 1

##### Benchmarking on simulated data

Sequencing data was simulated for ~200,000 human exons. For testing purposes, we artificially introduced an unrealistic high mutation rate of one INDEL per exon. INDEL sizes were randomly selected in the range [1,100] according to a lognormal distribution with mean $\mu$=1 and standard deviation $\sigma$=10. 100bp perfect reads were artificially generated using realistic coverage distribution based on probe patterns (30X average coverage). The reads were then aligned with both BWA-aln and BWA-mem [7] to the reference human genome hg19 using default parameters. Note that here we opted for a simple simulation model that, clearly, does not represent all of the complexities of a real exome project, e.g., potential cloning bias, sequencing errors, homopolymer runs, mutation hotspots, etc., but it is chosen only for the purpose of testing and comparing the best-case relative power of different pipelines to detect INDELs of different sizes under controlled conditions.

The figure below shows the size distribution of INDELs detected by seven different pipelines (Scalpel, SOAPindel [1], HaplotypeCaller, SAMtools [2], UnifiedGenotyper [3], FreeBayes [4], Platypus [5]) when applied to the simulated exome data. In accordance with previously published results, we report reduced power for standard mapping pipelines to detect large INDELs (specifically insertions): SAMtools, UnifiedGenotyper (GATK), and FreeBayes have limited power to discover deletions larger than 20bp and this trend is more evident for large insertions. The table shows the same trend: SAMtools, UnifiedGenotyper (GATK), and FreeBayes have lower sensitivity compared to the other tools. Surprisingly Platypus, although it is an assembly-based method, shows also limited power to detect longer INDELs and a significantly higher false discovery rate compared to Scalpel, SOAPindel, and HaplotypeCaller. Conversely, methods that use sequence assembly techniques are able to detect mutations over the full spectrum of the "true" distribution. Based on this analysis on simulated exome data, Scalpel, SOAPindel, and HaplotypeCaller have the best sensitivity, although Scalpel has the highest specificity. However, various simplifying assumptions used in the simulation may explain why simulated data appear somewhat easier to analyze, and the performance of these tools changes dramatically when applied to real exome capture data, as shown in the main text of the paper.

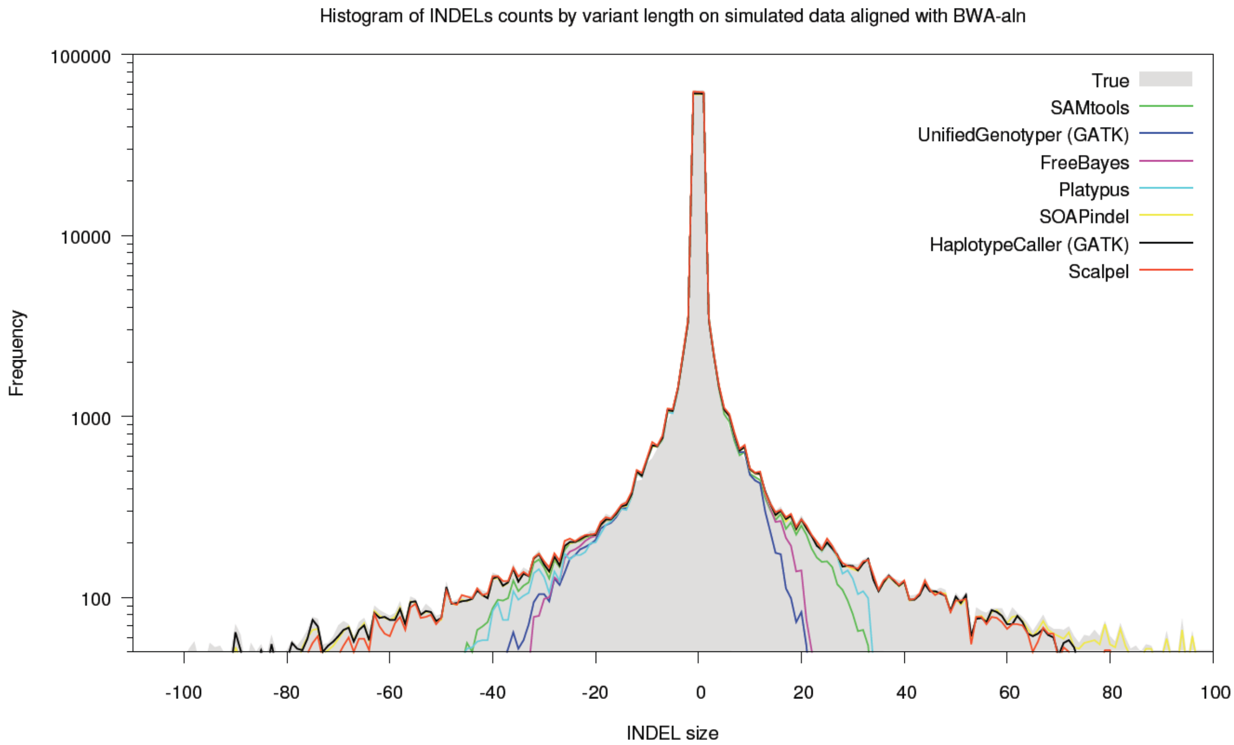


**INDELs size distribution of different pipelines on simulated exome data aligned with BWA-aln.** The true distribution of INDEL sizes is colored in grey. Standard mapping approaches ("SAMtools", "UnifiedGenotyper", "FreeBayes") show reduced power to detect longer mutations compared to assembly based pipelines ("Scalpel", "SOAPindel", and "HaplotypeCaller").

**Sensitivity and False Discovery Rate (FDR) for simulated data aligned with BWA-aln.** The first two columns consider all INDELs, whereas the last two just consider larger INDELs at least 5bp in size.

| Tool | Sensitivity (%) (all) | FDR (%) (all) | Sensitivity (%) (size > 5bp) | FDR (%) (size > 5bp) |
|---|---|---|---|---|
| Scalpel | 92.9 | 0.026 | 85.2 | 0.06 |
| SOAPindel | 93.3 | 0.070 | 89.7 | 0.09 |
| HaplotypeCaller | 93.7 | 0.110 | 87.8 | 0.49 |
| UnifiedGenotyper | 86.8 | 0.220 | 48.1 | 0.36 |
| SAMtools | 88.1 | 0.640 | 59.5 | 0.38 |
| FreeBayes | 85.8 | 0.212 | 44.3 | 0.07 |
| Platypus | 88.5 | 0.777 | 61.2 | 1.71 |

More recently a new version of BWA has been released (BWA-mem [6]) which is designed to be more sensitive to detecting longer INDELs. However, due to its new seeding algorithm, BWA-mem is likely to achieve higher sensitivity only for longer reads (>>100bp) and might not significantly improve the results for our simulated 100bp read dataset. We investigated if there was any improvement by aligning the simulated reads with the most recent version of BWA-mem. The Figure and the Table below show that for these data there is only a marginal improvement across all the tools in terms of sensitivity. The same trend of reduced power for standard mapping methods is clearly confirmed by the results. Unfortunately SOAPindel is not compatible with the BAM file generated by BWA-mem and it could not be included in the analysis (personal communications with BGI). The minor improvement in sensitivity is also confirmed by the alignment statistics for BWA-aln and BWA-mem where there is only a marginal increase in the number of mapped reads around INDELs. We used QualiMap [7] to compute the alignment statistics.

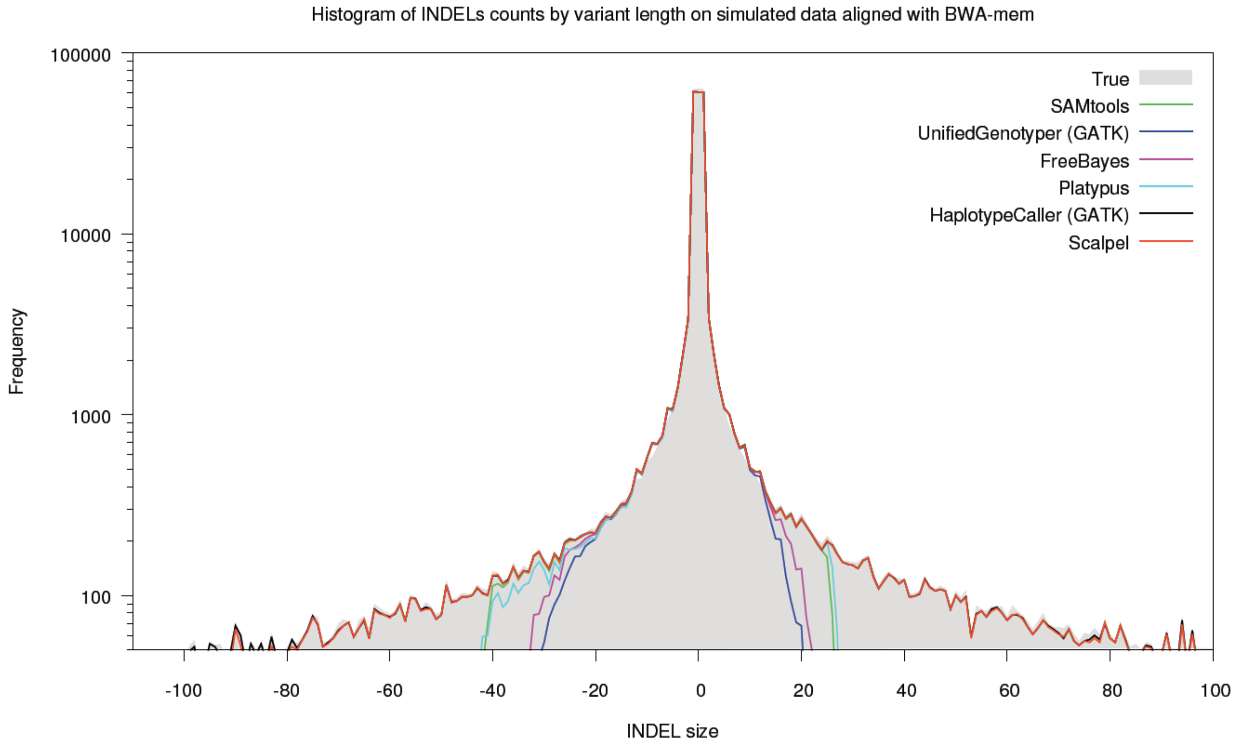


**INDELs size distribution of different pipelines on simulated exome data aligned with BWA-mem.** The true distribution of INDEL sizes is colored in grey. Standard mapping approaches ("SAMtools", "UnifiedGenotyper", "FreeBayes") show reduced power to detect longer mutations compared to assembly based pipelines ("Scalpel", "SOAPindel", and "HaplotypeCaller").

**Sensitivity and False Discovery Rate (FDR) for simulated data aligned with BWA-mem.** The first two columns consider all INDELs, whereas the last two just consider larger INDELs at least 5bp in size.

| Tool | Sensitivity (%) (all) | FDR (%) (all) | Sensitivity (%) (size > 5bp) | FDR (%) (size > 5bp) |
|---|---|---|---|---|
| Scalpel | 94.3 | 0.027 | 91.2 | 0.06 |
| SOAPindel | n.a. | n.a. | n.a. | n.a. |
| HaplotypeCaller | 94.4 | 0.068 | 91.8 | 0.17 |
| UnifiedGenotyper | 86.3 | 0.026 | 45.9 | 0.06 |
| SAMtools | 88.2 | 0.149 | 58.9 | 0.51 |
| FreeBayes | 86.9 | 0.013 | 49.8 | 0.04 |
| Platypus | 88.1 | 0.196 | 57.5 | 1.27 |

**Alignment statistics.** Comparison between BWA-aln and BWA-mem.

| | BWA-aln | BWA-mem |
|---|---|---|
| Number of reads | 50,549,624 | 50,659,783* |
| Mapped reads | 50,208,487 / 99.33% | 50,608,913 / 99.9% |
| Paired reads | 50,208,487 / 99.33% | 50,608,913 / 99.9% |
| Mapped reads, singletons | 57,769 / 0.11% | 15,362 / 0.03% |
| Read min/max/mean length | 45 / 100 / 100 | 30 / 100 / 99.86 |
| Clipped reads | 862,727 / 1.71% | 1,230,432 / 2.43% |
| Duplication rate | 39.23% | 39.36% |
| Total reads with indels | 6,458,280 | 6,706,818 |
| Insertions | 3,192,230 | 3,295,515 |
| Deletions | 3,266,050 | 3,411,303 |
| Homopolymer indels | 29.81% | 30.11% |

* Note that the total number of reads reported by flagstat is slightly higher for BWA-mem. This is due to the fact that BWA-mem marked a small portion of the reads as secondary alignment,. These reads were counted twice by QualiMap.

#### Supplementary Note 2

##### Comparison to RepeatSeq and lobSTR on simulated data

RepeatSeq [8] and lobSTR [9] are two recent computational tools specifically designed to detect INDELs in short repeat structures. In particular, RepeatSeq is designed to determine genotypes for microsatellite repeats in high-throughput sequencing data, while lobSTR is a tool for profiling Short Tandem Repeats (STRs) from high throughput sequencing data. As such, neither RepeatSeq nor lobSTR is a general-purpose INDEL caller for exome capture data, but they work only on selected regions of the human genome where these repeat structures are identified. For example lobSTR requires the input of a custom-made index file describing the STR loci. Out of 1638523 regions from the lobSTR index file, only 5436 are within the exonic region defined by our BED file used to generate the simulated data. Also, as expected, only a small fraction of these regions contain a mutation exactly within a STR sequence.

The tables below show the results of the comparative analysis between Scalpel, RepeatSeq and lobSTR. RepeatSeq and lobSTR show reduced sensitivity compared to Scalpel. This is particularly evident for longer INDELs (>5pb), where RepeatSeq and lobSTR demonstrate inferior power. This result is explained by the fact that both RepeatSeq and lobSTR are tools based on standard mapping approaches and, in agreement with the results reported in Supplementary Note 1, they have reduced power to detect large mutations.

| Tool | # of INDELS | True Positives | # of INDELS (>5bp) | True Positives (>5bp) |
|---|---|---|---|---|
| RepeatSeq | 476 | 472 | 36 | 36 |
| Scalpel | 656 | 656 | 164 | 164 |

| Tool | # of INDELS | True Positives | # of INDELS (>5bp) | True Positives (>5bp) |
|---|---|---|---|---|
| lobSTR | 431 | 370 | 56 | 22 |
| Scalpel | 662 | 662 | 166 | 166 |

#### Supplementary Note 3

##### Comparison between Scalpel and GATK (v2.4-3, v2.8, v3.0) on K8101-49685

The GATK pipeline is under active development. New versions of the software were released subsequent to our initial MiSeq experimental validations (see the Results and Online Methods sections). In this section, we compare two recent versions of the GATK (v2.8 and v3.0) with the published version that was used for our comparisons in the main text of the paper (v2.4-3). Specifically, in the figures below, we compare the INDEL size distribution of HaplotypeCaller from all 3 versions of the GATK package (v2.4-3, v2.8 and v3.0) using different filtering criteria. In the case of single exome studies, the GATK support group does not recommend using Variant Quality Score Recalibration (VQSR) because it is unlikely that the modeling step will perform well on a limited number of variants. Instead they advise using a hard filtering approach ("QD < 2.0 || FS > 200.0”). The three plots below show the comparison between the “raw” INDELs and the “hard-filtered” INDELs for the three versions of HaplotypeCaller. It is evident from the distribution of INDELs of v2.4-3 that hard filtering is a very aggressive strategy that drastically reduces the power of HaplotypeCaller to detect INDELs greater than 20bp long. This type of filtering seems to reduce HaplotypeCaller to the same power of the GATK’s UnifiedGenotyper, so we did not use this filtering step in our experimental comparison on sample K8101-49685. GATK’s HaplotypeCaller v3.0 instead does not show any significant difference in size distribution when the hard filter is applied, which is consistent with the GATK v3.0 release notes.

This comparison also shows that the new versions of HaplotypeCaller have significantly reduced the bias towards deletions, reported previously for GATK v2.4-3. Since most of the false-positive mutations reported by HaplotypeCaller v2.4-3 were large deletions, we devised additional experiments to evaluate the performance of GATK v3.0. Specifically, we randomly selected 200 INDELs specific to HaplotypeCaller v3.0 and 200 specific to Scalpel for targeted MiSeq validation (see the Online Methods section for the validation protocol).

Note this selection was blinded to any previous validation analysis, making it a fair, random selection from both algorithms. 215 out of the 400 INDELs were covered by more than 1000 reads in previous data sets, which includes data generated by the MiSeq validation experiment performed on the initial 1000 validation targets (see the Results and Online Methods sections) as well as data generated and reported in O'Rawe *et al.* 2013 [10]. We performed targeted MiSeq re-sequencing on the remaining 185 INDELs where data were not already available. 58 out of the remaining 185 were INDELs specific to Scalpel and 127 were INDELs specific to HaplotypeCaller v3.0. 170 out of the 185 INDELs were successfully sequenced and covered by more than 1000 reads, with an average coverage of 165142. The results of these re-sequencing experiments are reported in the two tables below. Under both the position-based and exact-match comparison strategies, Scalpel outperforms HaplotypeCaller v3.0. In particular, Scalpel shows a significantly higher validation rate for longer INDELs (size >5bp) as was previously observed with the older versions. We applaud the GATK developers for their improved algorithm, and expect them, along with us, to make further refinements over time as our understanding of the mutation mechanisms and error models improve.

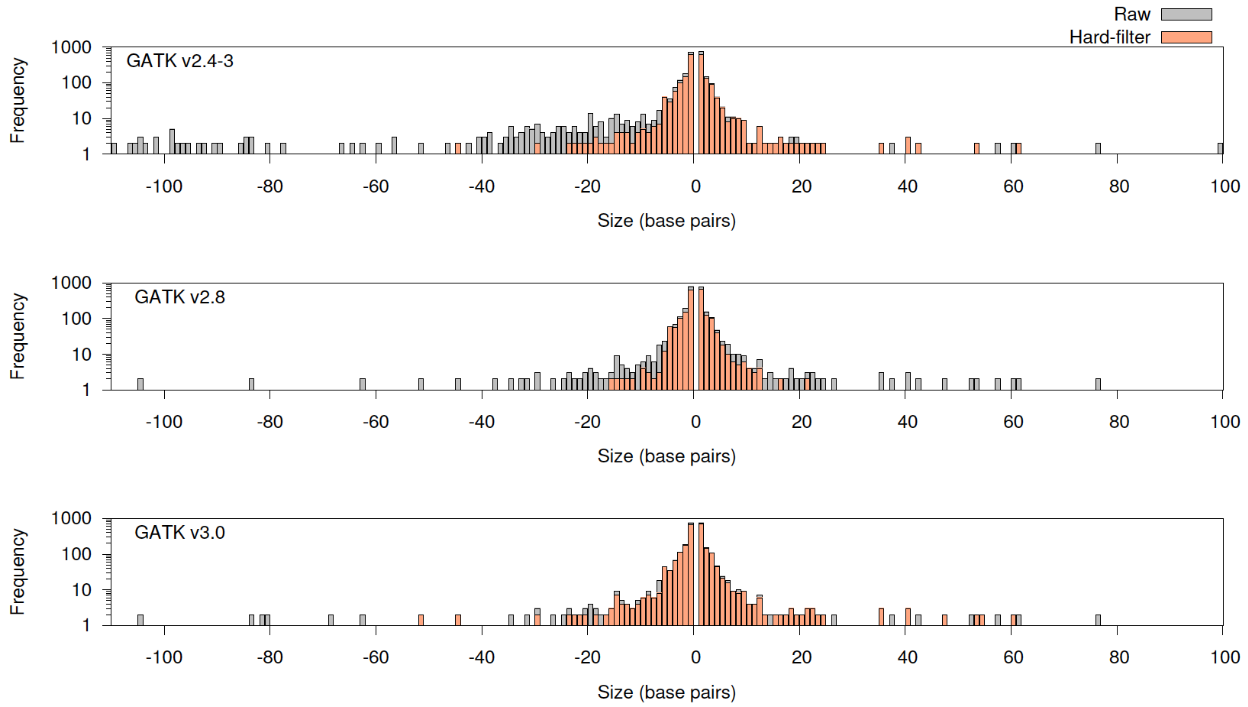


**HaplotypeCaller comparison using different filtering criteria.** Comparison between raw and hard-filtered INDELs for HaplotypeCaller in GATK v2.4-3, v2.8 and and v3.0.

**Validation rate for Scalpel and HaplotypeCaller (v3.0) using position-based comparison.**

| Tool | Valid | Invalid | PPV | Valid (>5bp) | Invalid (>5bp) | PPV (>5bp) |
|---|---|---|---|---|---|---|
| Scalpel | 146 | 50 | 75% | 26 | 4 | 86% |
| HapCall(v3.0) | 86 | 103 | 45% | 5 | 13 | 27% |

**Validation rate for Scalpel and HaplotypeCaller (v3.0) using exact-match comparison.**

| Tool | Valid | Invalid | PPV | Valid (>5bp) | Invalid (>5bp) | PPV (>5bp) |
|---|---|---|---|---|---|---|
| Scalpel | 82 | 114 | 42% | 23 | 7 | 76% |
| HapCall(v3.0) | 65 | 124 | 34% | 5 | 13 | 27% |

#### Supplementary Note 4

##### Characterization of validation rate

Among the different quality metrics computed by Scalpel, the chi-squared *k*-mer statistic ($\chi^2$) and the coverage ratio are the most informative scores. In this section we characterize the False-Discovery Rate (FDR) of Scalpel in real experiments as a function of both the chi-square statistics and coverage. In particular in the figures below we analyzed a total of 614 INDELs detected by Scalpel and validated by re-sequencing on the single-exome study (ID: K8101-49685). Specifically, given a fixed chi-square threshold T, we computed the FDR for all the mutations with chi-square score <T (x-axis). The corresponding FDR value is reported on the y-axes. Higher chi-square thresholds produce more errors but the FDR value stabilizes around 25%, in agreement with the experimental results reported in the main text of the manuscript. However, different trends are reported for mutations within microsatellites: even at low chi-square score the FDR for mutations within microsatellites is higher than the non-microsatellite mutations. This analysis also demonstrates the additional challenges faced in INDEL detection within microsatellites. Finally, different curves are plotted for subsets of mutations with predefined maximum coverage. As expected, mutations with higher coverage more frequently pass validation (lower FDR).

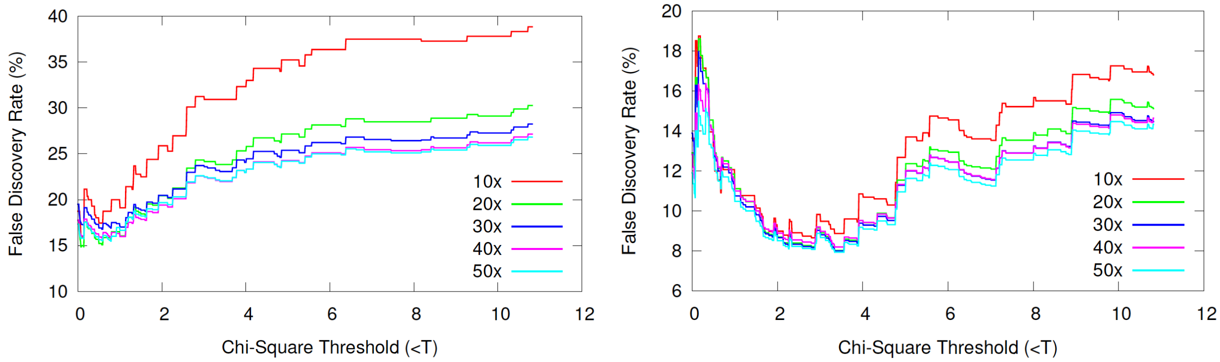


**Characterization of False Discovery Rate as a function of the chi-squared score and coverage.** The analysis is reported for mutations outside of microsatellites (left plot) and within microsatellites (right plot). Different curves are plotted for subsets of mutations with predefined maximum coverage.

We also analyzed the direct relationship between the chi-square score and coverage for the 614 mutations. The scattered plot below shows the same trend where mutations that did not pass validation (red dots) are generally characterized by lower coverage. Finally it is important to note that, since these results are based on a very small subset of only 614 mutations, there is a clear residual noise in both the previous FDR curves and the scattered plot blow. However the sample is large enough to start identifying the major trends and relationships between the validation rate and the characteristics of the data (such as chi-square score and coverage).

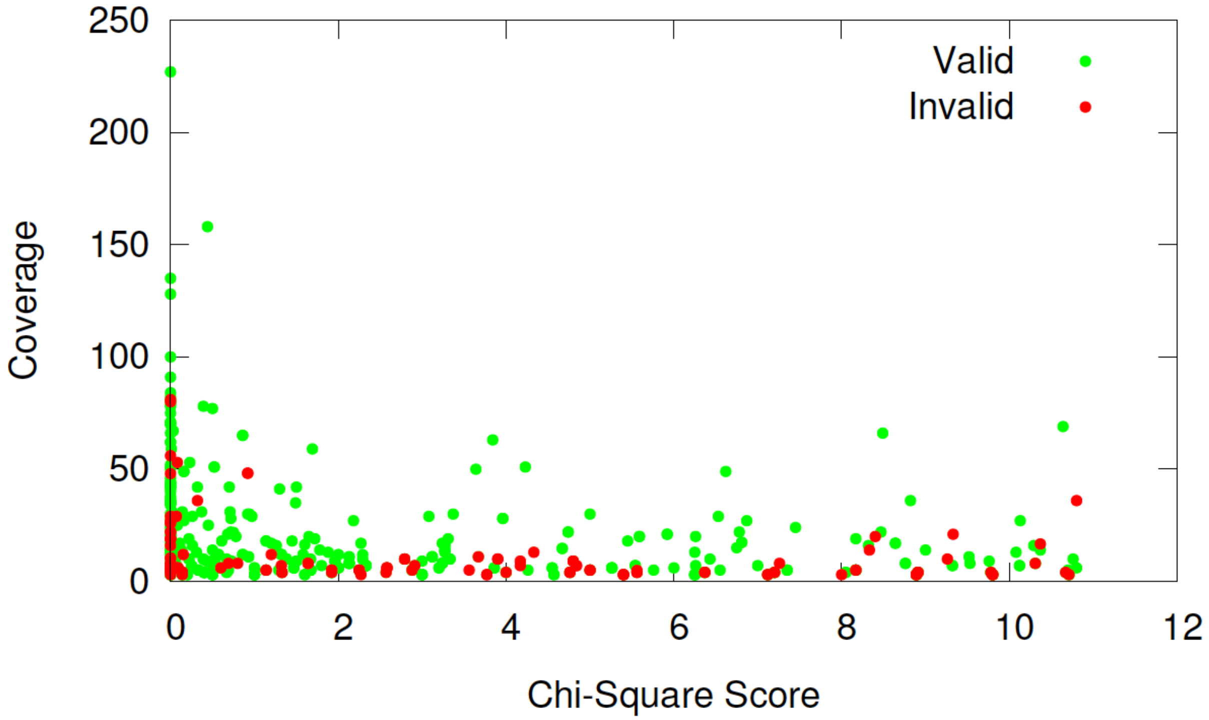

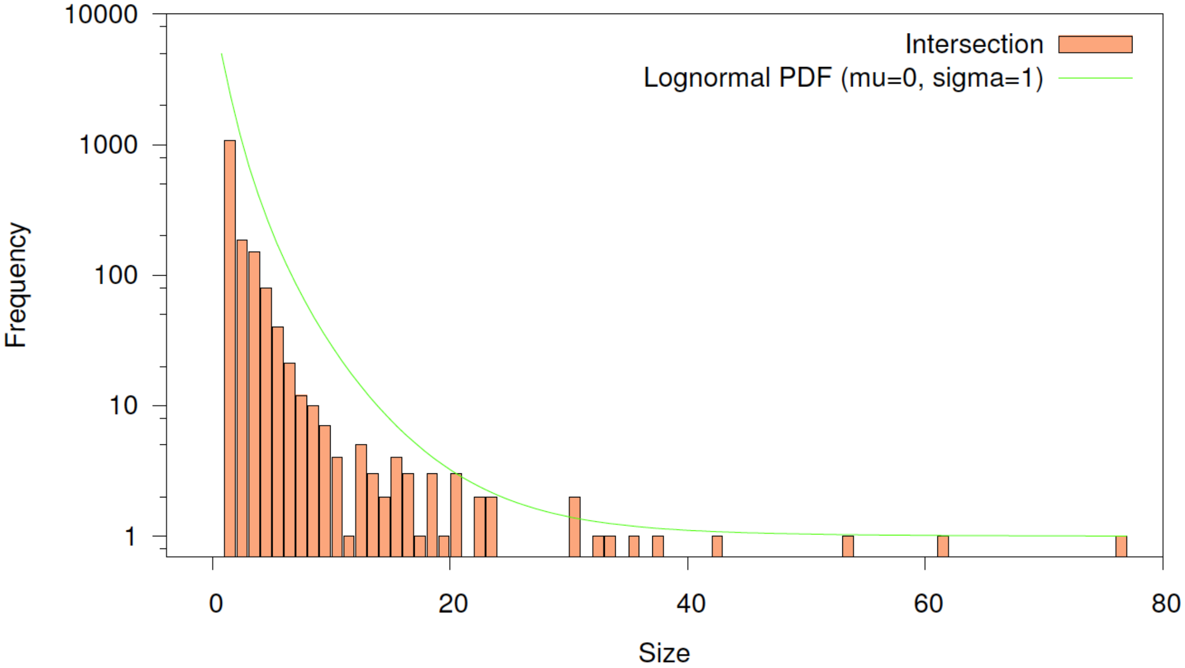


**Supplementary Figure 1. Size distribution of INDELs detected by all pipelines ("Intersection").** The sizes of INDEL belonging to the intersection among the three pipelines follow a lognormal distribution.

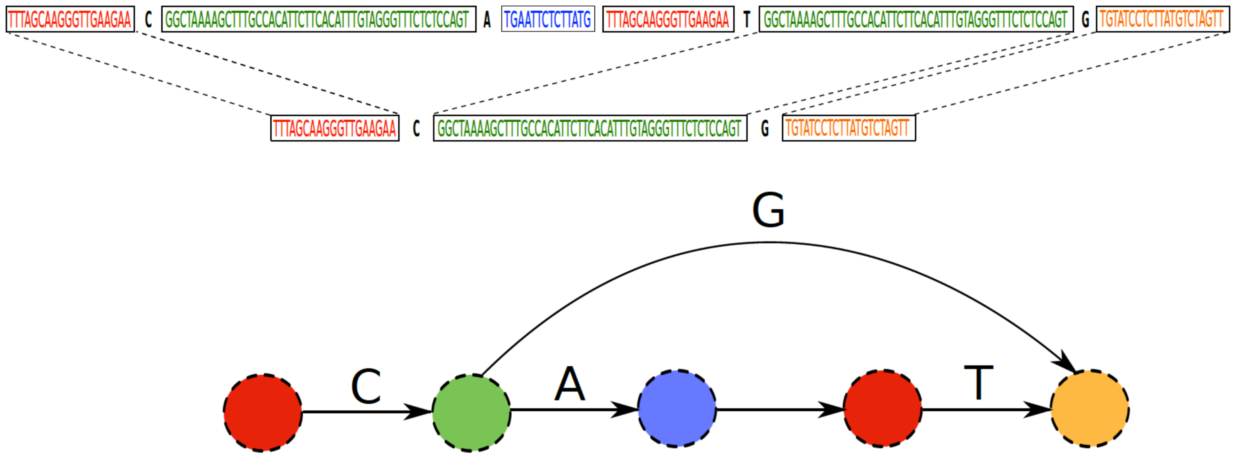


**Supplementary Figure 2. Example of false-bubble induced by a near perfect repeat sequence**. Near-perfect repeats are a major source of false positive variation calls when using a de Bruijn graph assembly strategy. This figure shows an example of near-perfect repeat that can be misinterpreted as a large deletion. The key point is that the beginning of this sequence is a nearly perfect 69bp repeat. There is just 1bp difference between the two copies that are 15bp apart. The sequence is segmented as 19-C-49-A-14-19-T-49-G-21 where 19 and 49 are 19bp and 49bp perfect repeats, separated by a 15bp unique sequence (A, C, T, G are the regular bases). Since the longest exact repeat is 49bp long, one would expect that using *k*-mer=55 should be large enough to correctly assemble this sequence. However, our data also contained reads with sequence 19-C-49-G, which suggests the presence of an 84bp deletion of the A-14-19-T-49 segment but in actuality is just a single base change. Since the de Bruijn graph is constructed using perfect matches of length *k*-1=54 (no mismatches allowed), the only way to connect all the 55-mers from these two sequences is to construct a false bubble jumping form the first copy of the near-perfect repeat to the second copy. When aligned to the reference, the sequence associated to the false branch will show a (false-positive) large deletion.

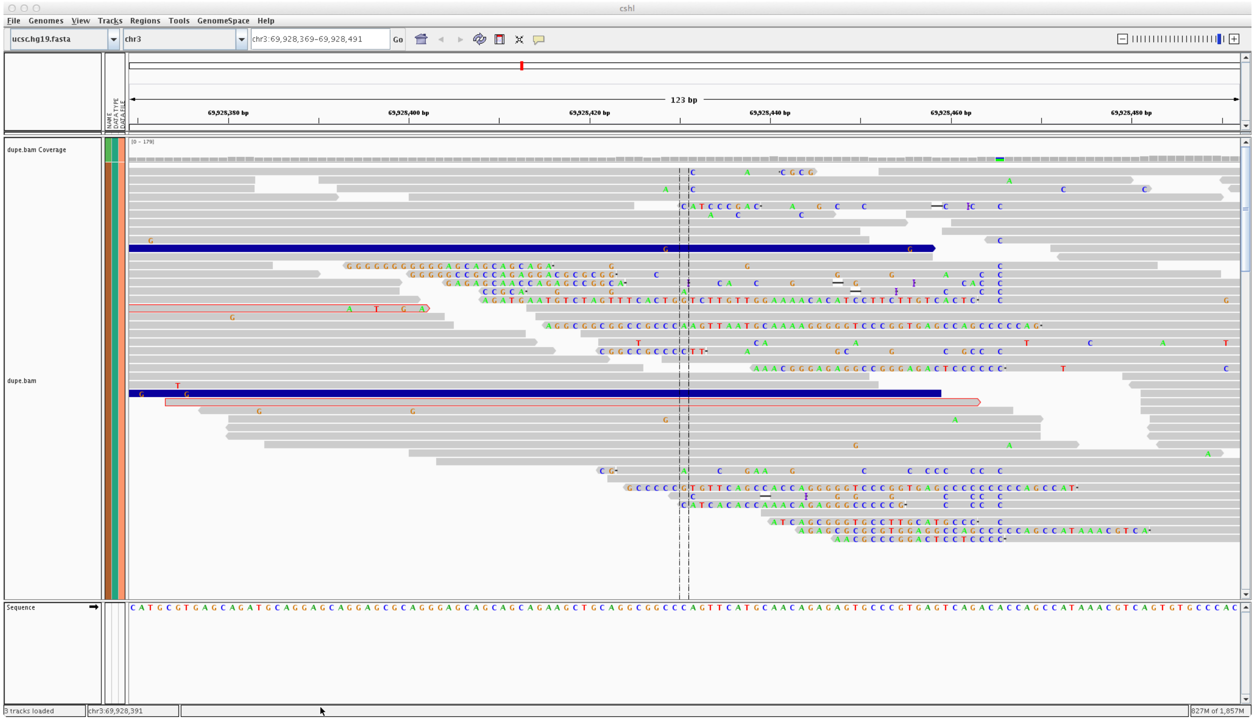

**Supplementary Figure 3. IGV snapshot of HaplotypeCaller false-positive deletion.** A 35 bp deletion was erroneously called by HaplotypeCaller at position 3:69928431. Reads aligned to this region show a cluster of highly variable soft-clipped sequences that are likely to be mapping artifacts.

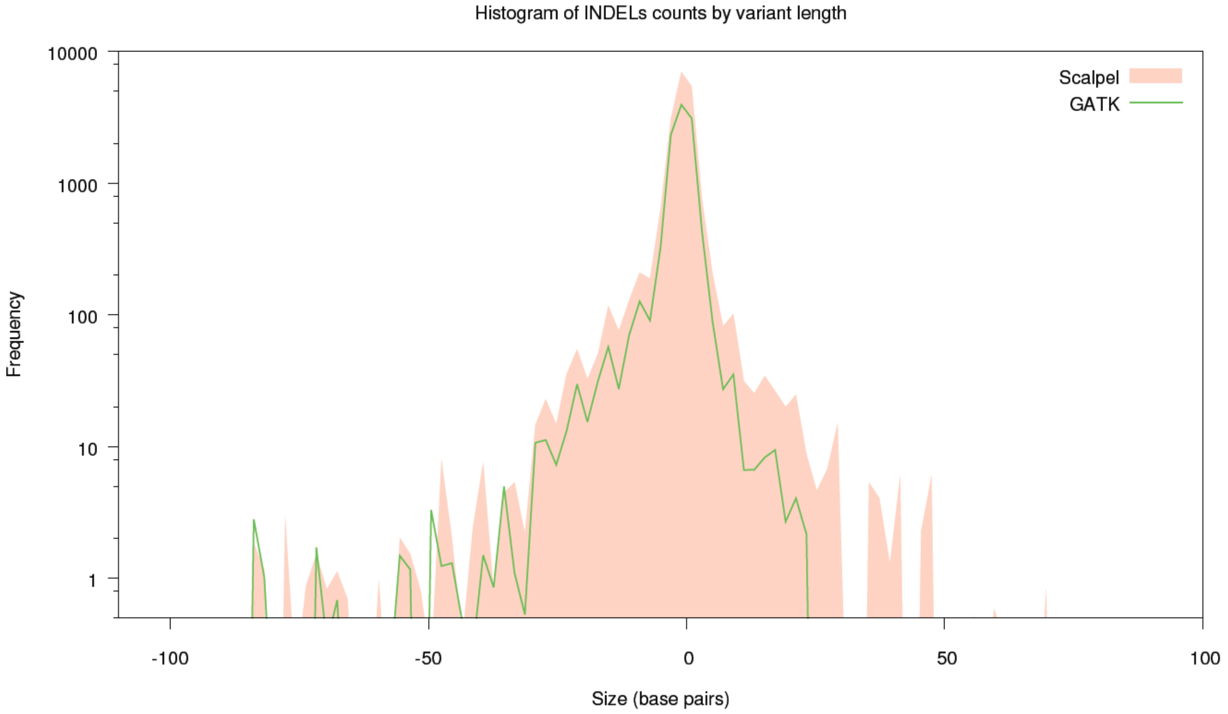


**Supplementary Figure 4. Comparison between Scalpel and GATK UnifiedGenotyper on 593 SSC families.** Scalpel demonstrates increased power to detect insertions.

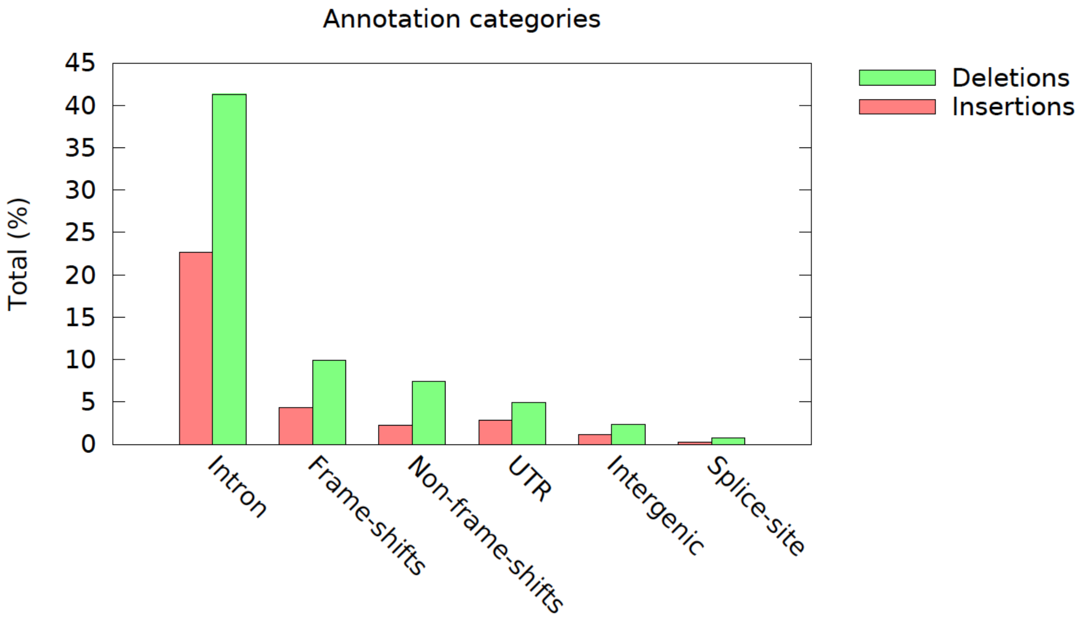


**Supplementary Figure 5. Abundance of INDELs for different annotation categories.**

**Supplementary Table 1. Number of insertions and deletions by annotation category.** DI-ratio is the ratio between the number of deletion and insertion for each category.

| Category | Insertions | Deletions | DI-ratio | Total |
|---|---|---|---|---|
| **Intron** | 6295 | 11490 | 1.82 | 17785 |
| **Intergenic** | 310 | 645 | 2.08 | 955 |
| **Frame-shift** | 1207 | 2758 | 2.28 | 3965 |
| **Non-frame-shift** | 622 | 2049 | 3.2 | 2671 |
| **UTR** | 790 | 1377 | 1.74 | 2167 |
| **Splice site** | 56 | 196 | 3.5 | 252 |
| **All** | 9280 | 18515 | 1.99 | 27795 |

#### Supplementary Note 5

##### Examples of de novo INDELs corrected by Scalpel

**Supplementary Figure 6** shows an example of de novo INDEL in the GNPTG gene that was initially detected as a 2bp deletion using the popular BWA+GATK UnifiedGenotyper pipeline. The same mutation was instead detected by Scalpel as a longer 33bp deletion and then confirmed by specific re-sequencing. IGV inspection shows a clear cluster of soft-clipped reads where BWA was unable to align the read end-to-end (**Supplementary Fig. 7**). Interestingly, the sequence composition of the soft-clipped sequences is consistently the same among all the soft-clipped reads. This is the typical signature of a candidate mutation that has been erroneously mapped. Targeted re-sequencing of the regions distinctly shows the deletion of 33 base pairs (**Supplementary Fig. 8**).

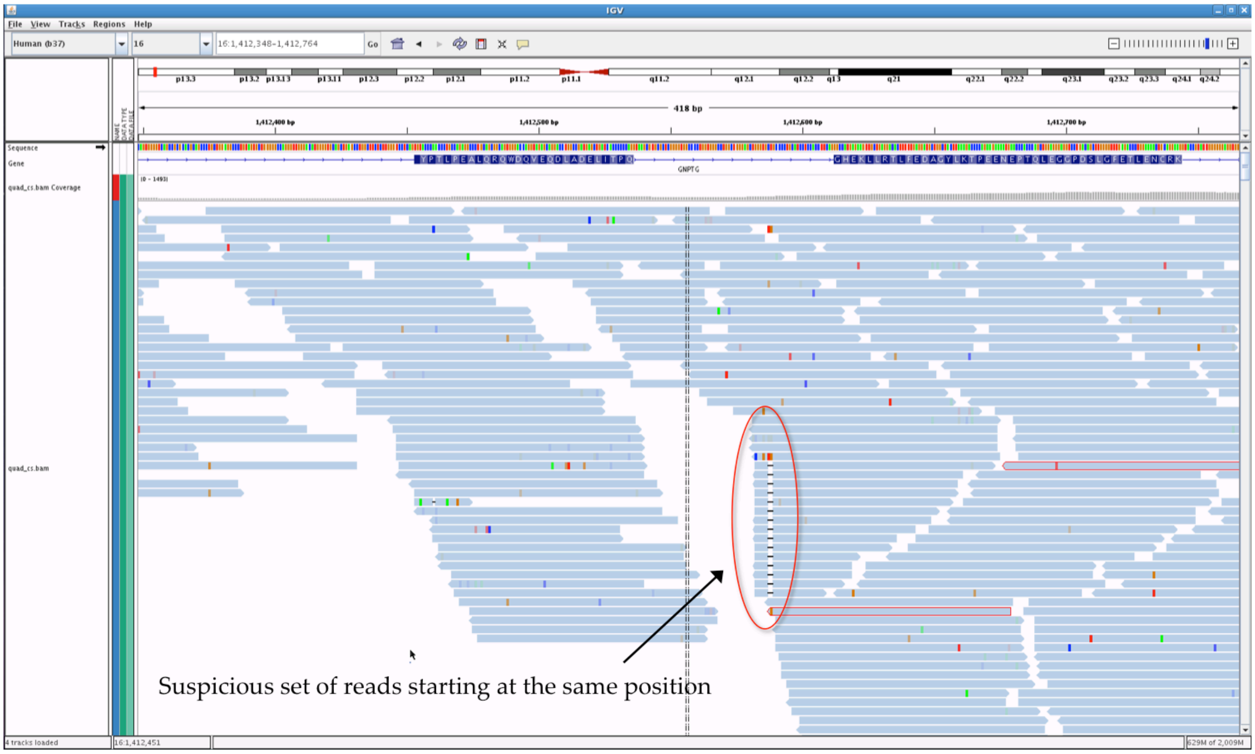


**Supplementary Figure 6. IGV inspection of a candidate 2bp deletion in the GNPTG gene.** The snapshot shows a suspicious set of reads starting at the same location.

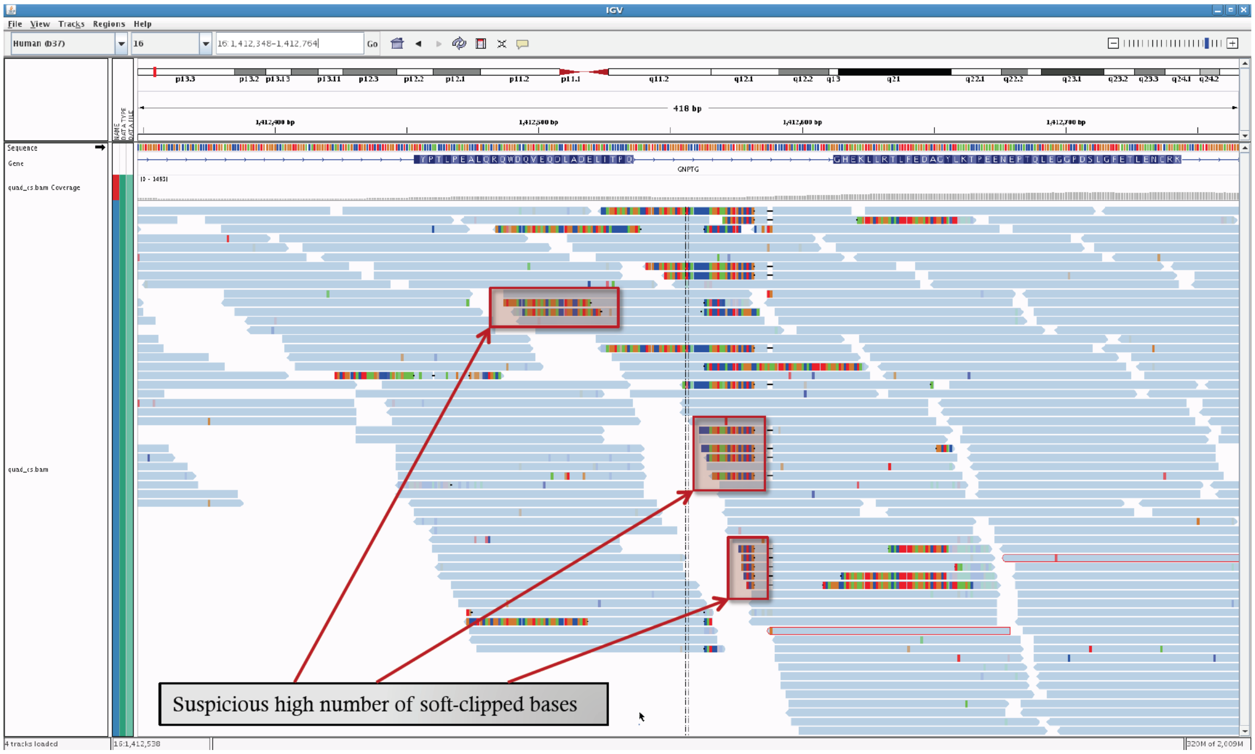


**Supplementary Figure 7. IGV inspection of the same 2bp deletion showing soft-clipped sequence.** IGV shows that the 2bp deletion from Supplementary Figure 6 is characterized by a suspicious cluster of soft-clipped reads**.**

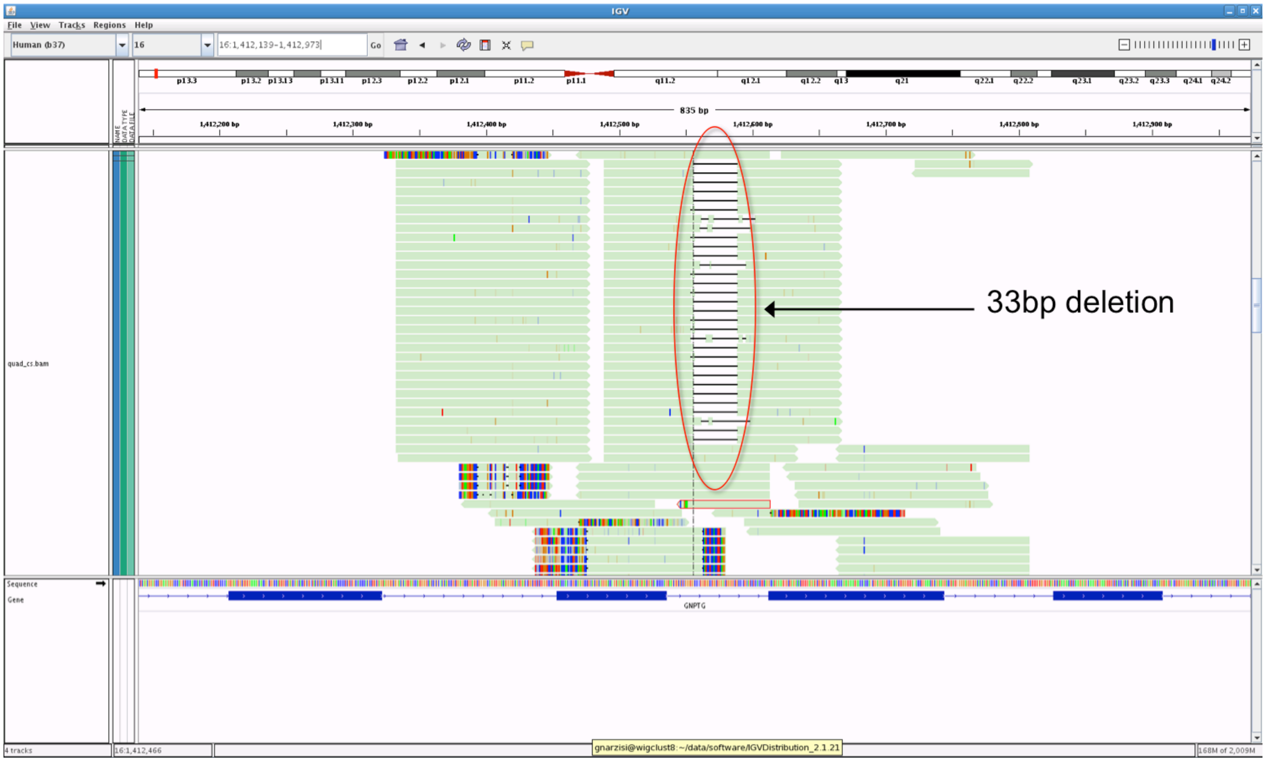


**Supplementary Figure 8. IGV inspection of a 33 bp deletion validated using longer reads.** Scalpel and specific re-sequencing confirm a large 33 bp deletion at the location (instead of a 2bp deletion).

In another region of chromosome 13, the GATK-based sequencing pipeline detects two different closely located deletions (1bp and 5bp) and, thus, they are flagged as LGD mutations (**Supplementary Fig. 9**). Scalpel instead detected only one deletion of 6 bp together with two adjacent SNPs detected in only one of the two haplotypes of the autistic child (family ID: 13578, location: 13:25280526), which was later confirmed by targeted re-sequencing (**Supplementary Fig. 10**).

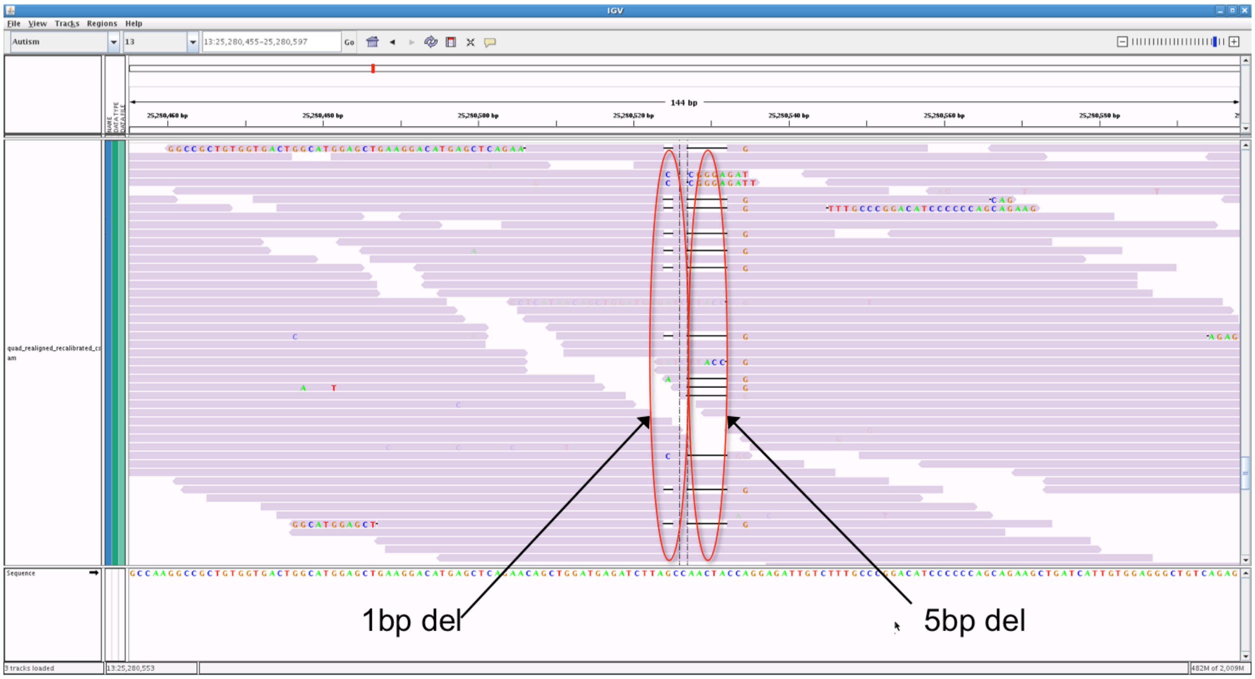


**Supplementary Figure 9. IGV inspection of two closely located deletions detected by the GATK-UnifiedGenotyper pipeline.**

```
Ref:      ...TCAGAACAGCTGGATGAGATCTTAGC|CAACTA|CCAGGAGATTGTCTTTGCCCGGA...
Mother:   ...TCAGAACAGCTGGATGAGATCTTAGC|CAACTA|CCAGGAGATTGTCTTTGCCCGGA...
Father:   ...TCAGAACAGCTGGATGAGATCTTAGC|CAACTA|CCAGGAGATTGTCTTTGCCCGGA...
Sib:      ...TCAGAACAGCTGGATGAGATCTTAGC|CAACTA|CCAGGAGATTGTCTTTGCCCGGA...
Aut(1):   ...TCAGAACAGCTGGATGAGATCTTAGC|CAACTA|CCAGGAGATTGTCTTTGCCCGGA...
Aut(2):   ...TCAGAACAGCTGGATGAGATCTTACC|------|CCGGGAGATTGTCTTTGCCCGGA...
```

**Supplementary Figure 10. Correct signature for the deletions of Supplementary Figure 8.** The correct signature as reported by Scalpel is a 6 bp deletion together with two adjacent SNPs detected in only one of the two haplotypes of the autistic child (family ID: 13578, location: 13:25280526).

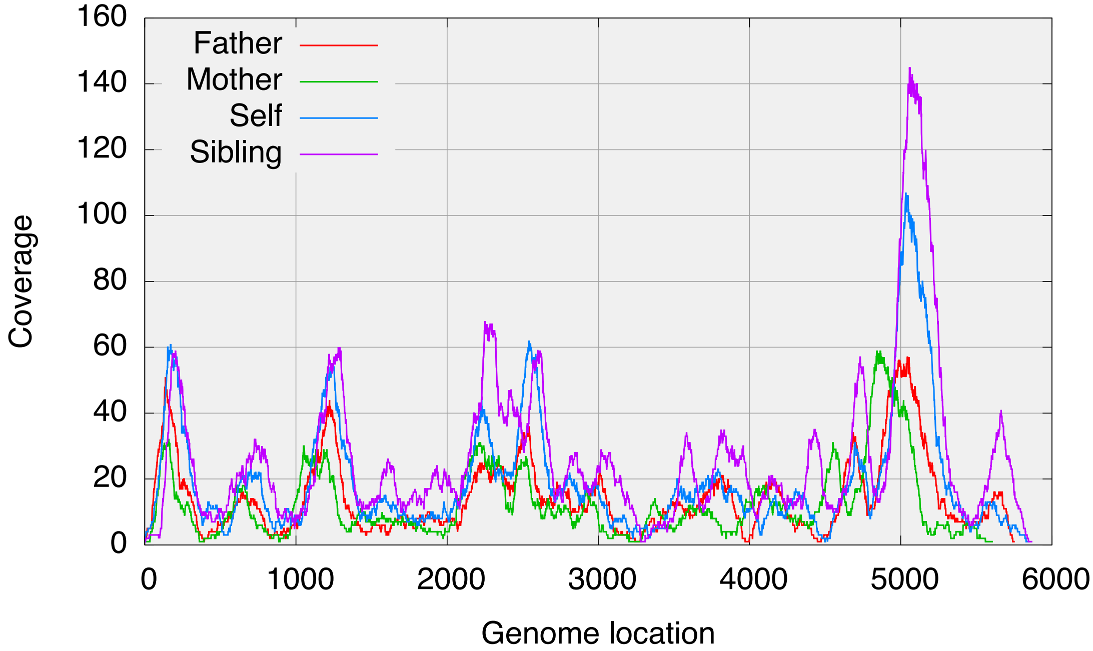


**Supplementary Figure 11. Read coverage across one long (6 kb) exon for one family of the SSC.** The highly variable distribution is clearly visible in the picture and it is consistent among the family members.

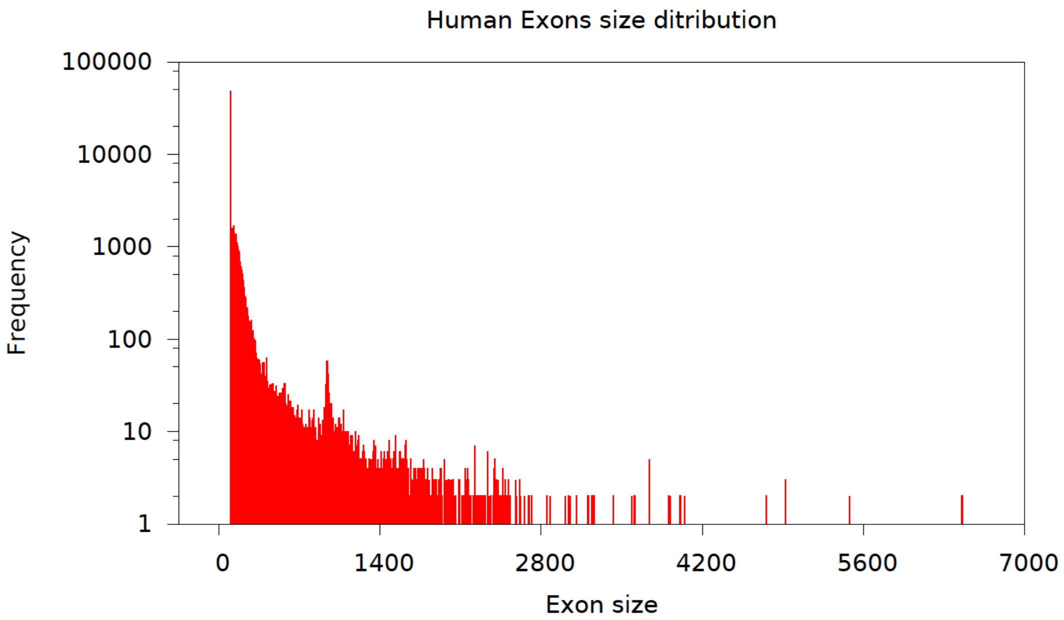


**Supplementary Figure 12. Size distribution of the human exon targets.** ~95% of the human exon-targets are shorter than 400bp

### Supplementary Note 6

#### Repeat composition analysis

Repeats are the most problematic sequences to assemble and the specificity of any INDEL detection method is very much correlated to its ability to detect and analyze repetitive sequences correctly. Although the exome sequence composition is generally assumed to be relatively simple compared to the rest of the human genome, 30% of exons have a perfect 10bp or larger repeat (**Supplementary Fig. 13**). But more interestingly the amount of *near-perfect* repeats increases substantially if more mismatches are permitted. **Supplementary Figure 13** shows the percent of locally repetitive exons as a function of different *k*-mer values and maximum number of mismatches. Each exon is analyzed to check for the presence of a repeat or near-perfect repeat in the same region. The y-axis reports the percentage of those blocks that have been found to contain a repeat of size *k*. Note that, given the generally low error rate of the Illumina technology, allowing 3-mismatches for a 10-mer (30% read accuracy) would not be realistic for a sequencing study. However, this repeat analysis must be examined in the context of performing sequence assembly using de Bruijn graph method. Since the input reads are decomposed into overlapping *k*-mers, the resulting assembly graph for a region with a near-perfect repeat can contain "false-bubbles" as the one depicted in Figure 2 of this Supplementary Information.

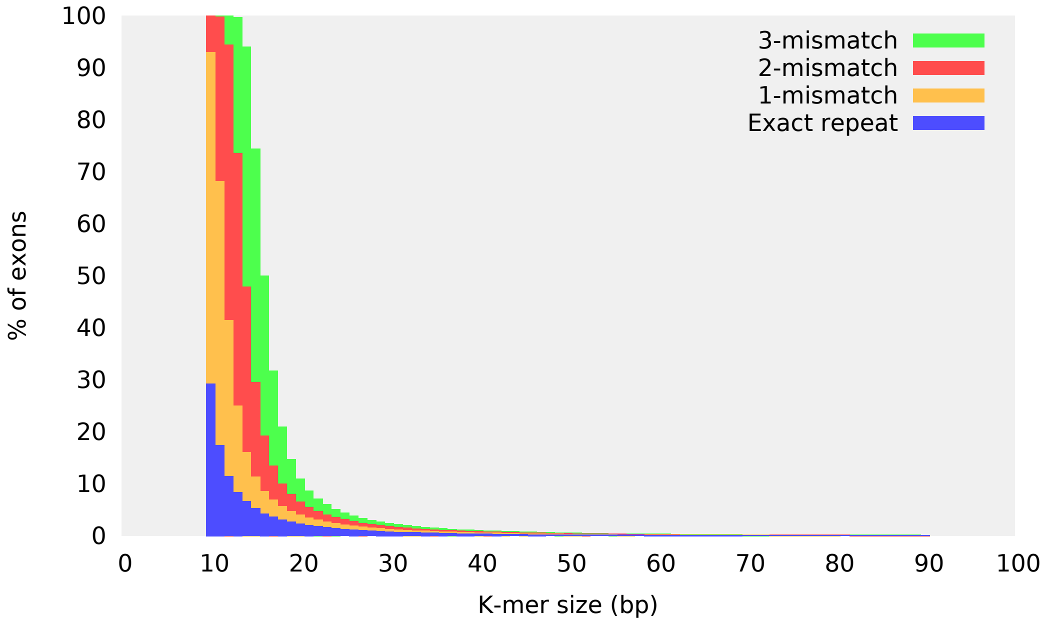


**Supplementary Figure 13. Repeat content of the Human Exome.** Repeat content distribution on the human exome targets as a function of the *k*-mer size. Each target is analyzed to check for the presence of a repeat or near-perfect repeat in the same region. The y-axis reports the percentage of those exons that have been found to contain a repeat of size *k*.

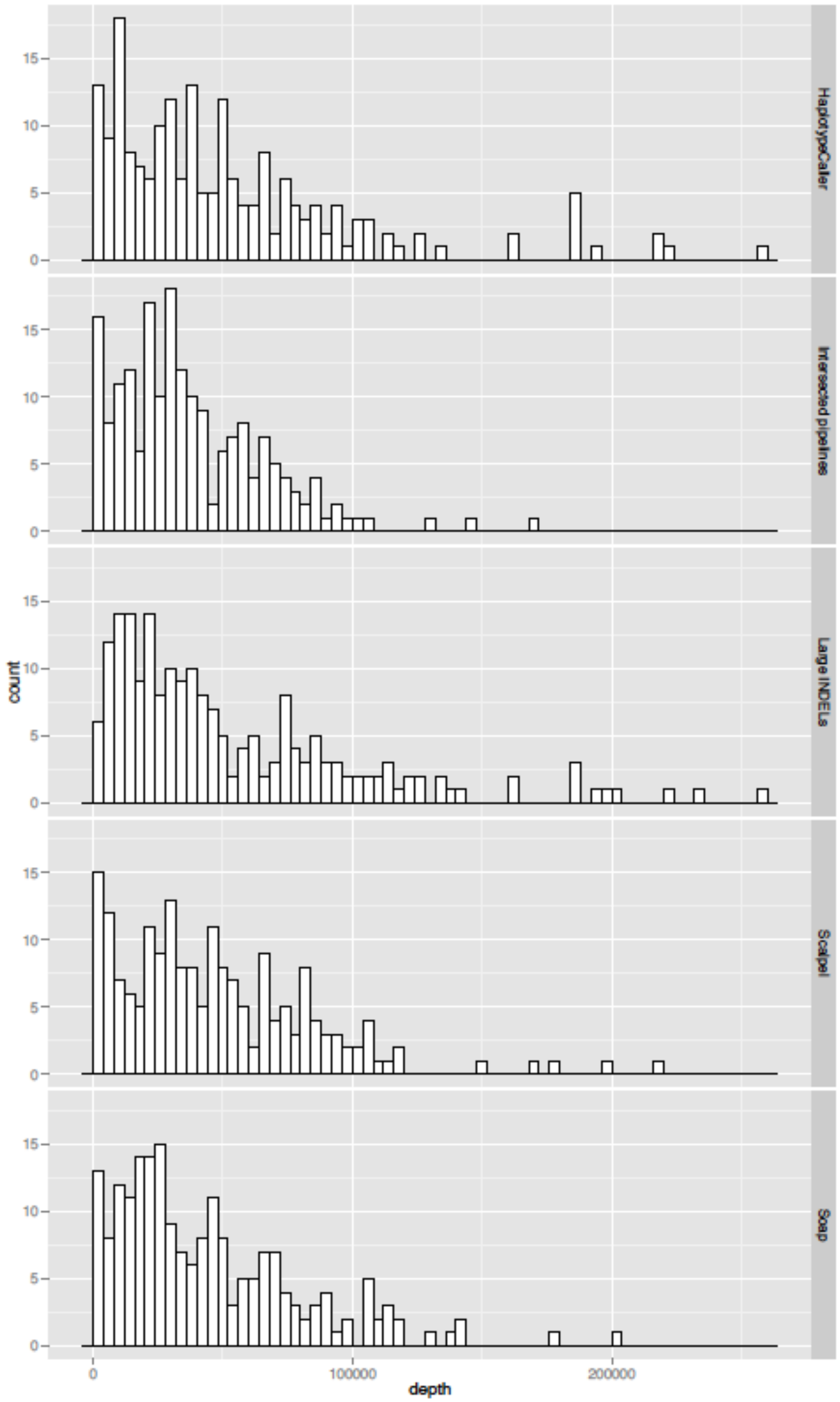


**Supplementary Figure 14. Histogram showing the coverage distribution for the INDELs validated with MiSeq.**
